# Social stress differentially affects the behavior, transcriptome, and signaling properties of the anterodorsal bed nucleus of the stria terminalis between male and female mice

**DOI:** 10.1101/2025.01.21.634142

**Authors:** Thomas J Degroat, Kevin M Moran, Katherine Denney, Sarah Paladino, Ali Yasrebi, Nadja Knox, Benjamin A Samuels, Troy A Roepke

## Abstract

Stress is comprised of systemic/physiological and processive/social factors, with social stressors requiring higher limbic processing. Sex is an important aspect of stress research as men and women show differing responses to stress and mood disorder development. We investigated how social stress affects the bed nucleus of the stria terminalis (BNST) and how sex may contribute. We used a chronic social instability stress (CSIS) paradigm to stress male and female mice for approximately 7 weeks. Afterwards, one cohort was used for avoidance behavior testing using the open field test, the elevated plus maze, the light/dark box emergence test, and the novelty suppressed feeding test. A second cohort was used for bulk RNA-sequencing of the anterodorsal (ad)BNST. A third cohort of CRH Cre+/Ai14 reporter mice were used for patch clamp electrophysiology in the adBNST. CSIS caused the females to be less avoidant, while the males became more avoidant. Low estrogen state in the females caused them to be less avoidant than in a high estrogen state. In the adBNST, we found major transcriptomic differences between the males and females. Males increased expression in more genes related to the RNA and protein processing whereas the females upregulated genes related to synaptic transmission. The transcriptome in the males is more sensitive to the stress than the females. Finally, CSIS caused adBNST CRH neurons to be hyperpolarized and have lower input resistance in males. In summary, social stress is differentially regulated between males and females, which may relate to the development of stress-related behavioral changes.

## 1. Introduction

When studying chronic stress, most research has used physical stressors, i.e. stressors that are direct affronts to physiology or homeostatic function. These types of stressors typically activate the fight-or-flight response to ready the body for immediate action to protect itself. This can be studied in animal models through multiple paradigms, including but not limited to chronic variable mild stress (Simpkiss and Devine, 2003), chronic restraint stress (Watanabe et al., 1992), and chronic foot shock (Kinn Rød et al., 2012). Researchers have modeled chronic stress outside of other environmental context through techniques such as chronic corticosterone administration (Woolley et al., 1990) or chronic corticotropin releasing hormone (CRH/F) administration (Krahn et al., 1990).

However, physical stressors are not the only type of stressors that organisms face. Stress can be further categorized into multiple different modalities, with a common delineation being stressors that require limbic processing, called processive stress, and stressors that do not, called systemic stress (Herman and Cullinan, 1997). Another axis is social versus physical stressors, which closely align with the ideas of systemic and processive stressors wherein social stress requires higher cognitive processing whereas many physical stressors do not. Instead, physical stressors directly activate the hypothalamic-pituitary-adrenal (HPA) axis and the sympathetic nervous system. For example, a meta-analysis of functional imaging of the human brain under physical versus social stress found that physical stress more preferentially activates the insula, cingulate cortex, and ventral striatum while social stress more preferentially activates the temporal gyrus and deactivates the dorsal striatum (Kogler et al., 2015). This suggests that physical stressors strongly activate the fight-or flight response while social stressors strongly activate mood regulation and motivation. We wanted to explore how physical stressors differ from social stressors in the central stress response by building on work we had previously done with a physiological chronic stress paradigm (Degroat et al., 2025, 2024) now with a socially-based stress paradigm.

Social stress has historically been difficult to model in animal studies. This is due to social stress typically requiring an understanding of complex concepts or social dynamics, as in situations such as financial stress, occupational stress, or relationship stress. One of the most common social stress paradigms in rodent models is social defeat stress, which relies on an aggressive male to dominate a subordinate or submissive mouse (Golden et al., 2011) However, the classic version of this paradigm is significantly limited in that it is not effective in females, as neither males nor females will fight an intruder female mouse under typical conditions (Harris et al., 2018). Though other techniques have been developed to allow for study of social defeat in females (Harris et al., 2018; Takahashi et al., 2017), these alternatives still require the primary stressor to be an attack from an aggressive male which can be seen as both physical and social stress. For this study, we opted to use a chronic social instability stress paradigm shown to be effective in both males and females (Yohn et al., 2019). This model may be more translatable to humans than other social stress paradigms like total social isolation.

The bed nucleus of the stria terminalis (BNST) is a limbic region of the brain that is selectively activated by long-context stressors (Perez-Villalba et al., 2005). As such, this region of the brain is important for chronic stress signaling and is often called the “extended amygdala.” However, the BNST is an extremely heterogeneous region, being able to be split into many subnuclei, each of which can have multiple different types of neurons (Beyeler and Dabrowska, 2020; Flanigan and Kash, 2022; Gegenhuber et al., 2022). We were specifically interested in the anterodorsal (ad) nuclei of the BNST as it has been shown to have connections to both the prefrontal cortex and the paraventricular hypothalamus, suggesting that is modulates both behavior and stress signaling (Xie et al., 2024). Additionally, the BNST has been shown to vary across sexes in morphology and expression of important signaling and hormone-related proteins (Allen and Gorski, 1990). Our own research found that not only is the transcriptome of the adBNST different between males and females, but the transcriptome is also differentially affected by physiological stress depending on sex (Degroat et al., 2025, 2024). This makes the adBNST an extremely compelling region for studying the sex variability in the development of stress-related mood disorders.

Overall, we sought to understand how social stressors uniquely and specifically affect the adBNST and how these effects may lead to behavioral changes. We hypothesized that social stress would result in an increase in avoidant behaviors, but that this increase would differ by sex and hormone status, as seen in our previous publication (Degroat et al., 2024). We also predicted that the adBNST transcriptome would include genes that are differentially expressed by sex and stress condition, with males being more sensitive to stress as previously shown (Degroat et al., 2025). Finally, we hypothesized that the electrophysiological properties such as the resting membrane potential, M-current, and excitatory postsynaptic current inputs of CRH neurons in the adBNST would be affected by chronic social stress exposure, particularly in males, mirroring effects of chronic variable mild stressors as previously observed (Degroat et al., 2025).

## 2. Materials & Methods

### 2.1 Animals

All procedures were conducted in accordance with National Institutes of Health standards, approved by the Rutgers Institutional Animal Care and Use Committee, and agree with ARRIVE’s guidelines for reporting animal research. Mice used for behavioral tests were C57BL/6J adult male and female mice bred in house. Mice used for RNA- sequencing cohorts were C57BL/6J adult male and female mice purchased from the Jackson Laboratory (Cat. # 000664). Mice were housed in a temperature and humidity-controlled room (22°C, 30-70% humidity), on a 12/12 h light/dark cycle, and provided food and water ad libitum. Control or stressed conditions started at 7-10 weeks of age. Due to the nature of the CSIS schedule, sample sizes were n=15 mice for each experimental group which allowed detection of effects with at least power = 0.8 at α = 0.05.

For electrophysiology, mice were cross-bred between an Ai14 parent and a corticotropin releasing hormone (CRH)- Cre parent such that offsprings had alleles for both Ai14 and CRH-Cre, resulting in exclusive expression of tdTomato in CRH-expressing cells. Both Ai14 and CRH-Cre breeding mice were purchased from Jackson Laboratories (Cat # 007914 & 012704, respectively).

### 2.2 Chronic Social Instability Stress Paradigm

The chronic social instability stress (CSIS) paradigm lasts for over 7 weeks and involves frequent changes in cage mates (Yohn et al., 2019). In this paradigm, every 3 days the mice are trio housed with novel cage mates, only allowing repeat cage conformations at most every 2 weeks to allow enough time for extinction of possible habituation to repeated cage mate exposure. For this, mice are taken out of their current cage, identities of the mice are verified, and their new home cage is determined. This is repeated for every mouse until all mice are with 2 novel cage mates. This frequent changing ensures that stable social hierarchies cannot be established. Control mice are also trio housed and have cages changed every 3 days to match CSIS groups, but control mice are never exposed to novel mice. Under these conditions, the only difference between the control and stress groups is the changing social dynamics.

### 2.3 Corticosterone Assay & Body Weight

Corticosterone (CORT) levels in the serum were analyzed as previously described (Degroat et al., 2024). Briefly, blood was collected at 3 timepoints in the behavior cohort. Blood was collected prior to stress via submandibular bleeds, after stress but before behavior via submandibular bleeds, and after behavior at termination via trunk blood after decapitation. CORT concentrations were then analyzed via a commercially available sandwich ELISA kit (#RTC002R; BioVendor). The lower limit for detection in this kit is 6.1 ng/ml with an intra-assay coefficient of variation between 5.9% and 8.9% and an inter-assay coefficient of variation between 7.2% and 7.5%. A logarithmic regression was then used to convert the absorbance to concentrations in ng/ml. A second analysis was done in terminal blood in which males were excluded and females were separated by estrus stage. Body weight (BW) was measured weekly and cumulative % BW change was calculated.

### 2.4 Behavior Assays

Behavioral assays were conducted over 8 days, alternating days between male and female testing so that mice were not affected by opposite-sex pheromones in the testing chambers. The tests conducted were the open field test (OFT), the elevated plus maze (EPM), the light/dark box emergence test (LDB), and the novelty suppressed feeding (NSF) test in that order. Vaginal cytology was performed on all females immediately after behavioral testing to monitor estrus stage. All tests were conducted as previously described (Degroat et al., 2024).

#### 2.4.1. Open Field Test

The OFT tests avoidant behaviors by testing a mouse’s willingness to explore an open arena which would be perceived as dangerous to a prey animal. The OFT is conducted in an opaque plexiglass box with an open top (40 cm long x 40 cm wide x 40 cm tall). The mouse is allowed to freely explore the arena for 10 minutes and entrances into and the amount of time spent in different zones is recorded. A mouse that spends more time in the perimeter and corner zones suggests a depressive-like increase in avoidant behaviors whereas a mouse that explores the center is less avoidant.

#### 2.4.2 Elevated Plus Maze

The EPM also tests avoidant behavior by giving the mouse a choice to explore either a perceived safe or a risky environment. EPM is conducted in an opaque plexiglass plus-shaped maze where two arms are enclosed by walls on three sides and the other two arms have no walls. Arms are 30cm long x 5 cm wide and connected at a 5 cm square center. The walls surrounding the closed arms are 15 cm tall with open tops. For this test, we analyzed the amount of time spent and the number of entries into the different arms. A mouse that spends more time in the closed arms is more avoidant than a mouse that explores the open arms.

#### 2.4.3 Light/Dark Box Emergence Test

The LDB test uses the same arena as the OFT, but an insert is placed inside to cover half the arena in darkness. This tests the mouse’s willingness to leave the perceived safety of dark areas and explore light areas. The insert is an opaque box (20 cm long x 40 cm wide x 40 cm tall) with a closed top and a hole at the bottom to serve as an entryway between light and dark areas. For this test, we measured the amount of time spent in each zone and the number of entries between zones. A mouse that spends more time in the dark is considered more avoidant.

#### 2.4.4 Novelty Suppressed Feeding Test

The NSF is the only test that does not strictly measure avoidance behavior. For this test, mice are fasted for 24 hours prior to two trials. For the first trial, mice are placed in a novel arena with a food pellet secured at the center. The mouse then must decide if they will stay in the safety of the edges or move to the center to obtain the pellet. The second trial is similar to the first but is conducted in their home cage instead of a novel arena. The NSF tests the interaction between avoidance and motivation. We analyze the latency to approach and eat the pellet in both arenas, the likelihood to eat in each arena, and the body weight changes due to fasting. A mouse that takes longer or does not eat the pellet at all is seen as more avoidant than one that quickly approaches the pellet.

### 2.5 RNA-Sequencing

For RNA-sequencing, mice were euthanized via rapid decapitation with ketamine. The brain was then dissected out and sliced for microdissection in which the anterodorsal (ad) BNST was isolated. These samples were placed in RNAlater (Invitrogen) solution for storage. Prior to RNA extraction, three biological replicates were pooled to account for interindividual differences resulting in an n=5 for each experimental group. RNA was extracted using the RNAqueous Micro Isolation kit (Invitrogen; AM1931). RNA concentration and integrity were then measured with an Agilent 2100 Bioanalyzer. All samples had RNA integrity numbers >8. Samples were then sent to the J.P. Sulzberger Columbia Genome Center at Columbia University (New York, NY) for library preparation and sequencing. Library preparation was performed with the TruSeq Stranded mRNA Library Prep Kit (Illumina) and pooled libraries were sequenced on the Illumina NovaSeq 6000 with 100 bp paired-end reads at a depth of 40 million reads. The resulting reads were quality trimmed with the FastX Toolkit (http://hannonlab.cshl.edu/fastx_toolkit/) with a minimum quality score of 20.0. They were then aligned to the mm10 reference genome using STAR and the NCBI RefSeq mm10 gene annotation. Reads were counted with FeatureCounts (Dobin et al., 2013; Liao et al., 2014). Data was then imported to R (Version 4.5.0) and differential expression was analyzed using DESeq2 (Version 1.46.0) with the following design parameters: (design ∼ sex + condition + sex:condition) (Love et al., 2014). Principle component analyses (PCA) were generated by PCAExplorer to check for outliers (Marini and Binder, 2019). A full list of all differentially expressed genes by comparison can be found in the supplemental materials. Gene ontology terms were analyzed with clusterProfiler (Wu et al., 2021).

### 2.6 Electrophysiology

Electrophysiology was conducted as previously described (Degroat et al., 2025). Briefly, mice were decapitated without anesthetization between ZT 16-18 (4 to 6 hours into the light cycle). The brain was then dissected out and sliced on a VT1000 S vibrating blade microtome (Leica) at 250 nm thickness in a high sucrose artificial cerebral spinal fluid (aCSF) that was prepared in house (containing in mmol: 208 sucrose, 2 KCl, 26 NaHCO3, 10 glucose, 1.25 NaH2PO4, 2 MgSO4, 1 MgCl2, 10 HEPES with a pH of 7.35 and osmolarity of 290–310 mOsm). Slices were then allowed to acclimate at room temperature in a bath of continuously oxygenated aCSF (containing in mmol: 124 NaCl, 5 KCl, 2.6 NaH2PO4, 2 MgCl2, 2 CaCl2, 26 NaHCO3, 10 glucose with a pH of 7.35 and osmolarity of 290-310). Once acclimated, individual slices were placed in the bath of the electrophysiology rig containing the same aCSF recipe as the acclimatization chamber and held at 35 . Individual cells were selected by fluorescence of the tdTomato reporter (a representative image of the fluorescing CRH+ neurons is available in fig. 7g). A borosilicate pipette backfilled with an internal solution (containing in mmol: 10 NaCl, 128 K-gluconate, 1 MgCl2, 10 HEPES, 1 ATP, 1.1 EGTA, and 0.25 GTP with a 7.35 pH and 290–300 mOsm) was used to patch onto the cell. Membrane values and standard protocol were recorded for every cell and throughout the duration of the patching to ensure the health of the cell. Data collection was done using 1550B Axon Instruments Digidata acquisition system, 700B Axon Instruments Multiclamp amplifier, and pCLAMP software (version 11.1) from Molecular Devices.

Recordings and analysis of the M-current were conducted as previously described (Degroat et al., 2025). Briefly, slices were first bathed in an aCSF solution that contained 1 μM tetrodotoxin (TTX) (Tocris, Cat. # 1069) and allowed to perfuse for 5 minutes before recording. Under voltage clamp, a standard deactivation protocol was used to measure K+ currents elicited during 500-ms voltage steps from 30 to 75 mV in 5-mV increments after a 300-ms prepulse to 20 mV. After this baseline recording, XE-991 (Tocris, Cat. # 2000), a selective inhibitor of KCNQ channels, was introduced to the bath at 40 μM and allowed to perfuse for 10 minutes. The same deactivation protocol was then recorded. M-current data was analyzed by Clampfit 11. The M-current is an inhibitory current that acts through voltage-gated K+ channels (Brown and Adams, 1980; Robbins, 2001). As one of the few currents that is active at physiologically relevant voltages, the M-current impacts the likelihood of a cell to fire an action potential (Marrion, 1997). In summation, a strong M-current suggests a cell that is less likely to fire, whereas a weak M- current suggests a cell that is more likely to fire. Representative traces of the M-current can be found in fig. 7h.

EPSCs were collected during a 5-minute recording under voltage clamp at −50 mV and analyzed on Easy Electrophysiology. EPSCs were detected using the template event detection feature using the deconvolution method with a theta of 4.5. Events were then averaged, and the number of events, average amplitude, and average area under the curve (AUC) for the EPSCs, were recorded. Representative traces of current recording for EPSCs can be found in fig 7c.

M-current tests had sample sizes of n = 20 for control males from ten mice, n = 18 for stressed males from nine mice, n = 13 for control females from six mice, and n = 13 for stressed females from six mice. There was one outlier for M-current removed from subsequent analyses. EPSC tests had sample sizes of n = 20 for control males from ten mice, n = 18 for stressed males from nine mice, n = 15 for control females from six mice, and n = 14 for stressed females from six mice. Females were tested only in diestrus to ensure consistent hormone profiles between mice; diestrus was confirmed by vaginal cytology the morning of the electrophysiology recording. Unless otherwise specified, all chemicals were purchased from Sigma Aldrich.

### 2.7 Statistics

All parameters were analyzed using Prism (GraphPad Prism, Version 10, Dotmatics) and used a p-value or adjusted p-value (false discovery rate, FDR) of <0.05 for significance unless otherwise specified. CORT assays and behavior were tested for outliers using the Grubbs test and then was analyzed by two-way ANOVA. BW was analyzed by a repeated-measures ANOVA. Behavior and CORT were first analyzed comparing treatment and sex. A second analysis followed that excluded males and separated females by estrus stage into proestrus and estrus (P/E) which represented a high estrogenic, hormonal flux state and metestrus and diestrus (D) which represented a low estrogenic, stable state. The resulting variables were then treatment and estrus stage. For the CORT levels, this was only done in terminal blood. Probability to eat in the novel arena and home arena were analyzed by Kaplan-Meier survival curves. RNA-Sequencing was analyzed via DESeq2 in R. A table of sample sizes and excluded data, including outliers and censored data, can be found in the supplemental materials.

Electrophysiology was analyzed using R v4.3.2 (R Core Team, 2014; Wickham et al., 2019). RMP, input resistance, and EPSC measures were analyzed with linear models, then tested post-hoc with Welch’s t-tests. M-current was analyzed with repeated measures two way ANOVA in Prism (Version 10). Results of linear models are presented as (b = beta coefficient ± standard error; n = number of observations in analysis; p = p value with α = 0.05, two-tailed). All post-hoc tests used Welch’s t-tests [mean ± standard deviation; t(df) t value with estimated degrees of freedom; p = p value with α = 0.05, two-tailed]. Effect size was calculated with (d = Cohen’s D).

## 3. Results

### 3.1 Body Weight & Corticosterone

The CSIS paradigm only significantly impacted body weight in the males used for RNA-sequencing. For the RNA- sequencing cohorts, control males gained 16.69 ± 0.6 % of their BW and stressed males gained 23.16 ± 1.78 % over the course of the 7-week paradigm (fig. 1a). This was a significant effect of treatment as the stressed males gained more weight than the controls (p = 0.0064, F (1, 28) = 8.704). Female controls gained 15.13% ± 1.2 % while stressed females gained 17.4% ± 1.06 % (fig. 1a). There were no significant treatment differences in female BWs. The behavior cohorts had a less drastic change in their BW, possibly due to the behavior cohorts being on average older than the RNA-Sequencing cohorts. The control males in the behavior cohort gained 6.61 ± 1.33 % while stressed males gained 8.46 ± 2.26 % (fig. 1b). Female controls gained 8.25 ± 0.96 % while stressed females gained 7.75 ± 1.98 % (fig. 1b). Neither the males nor females of the behavior cohorts significantly differed.

**Fig. 1.**
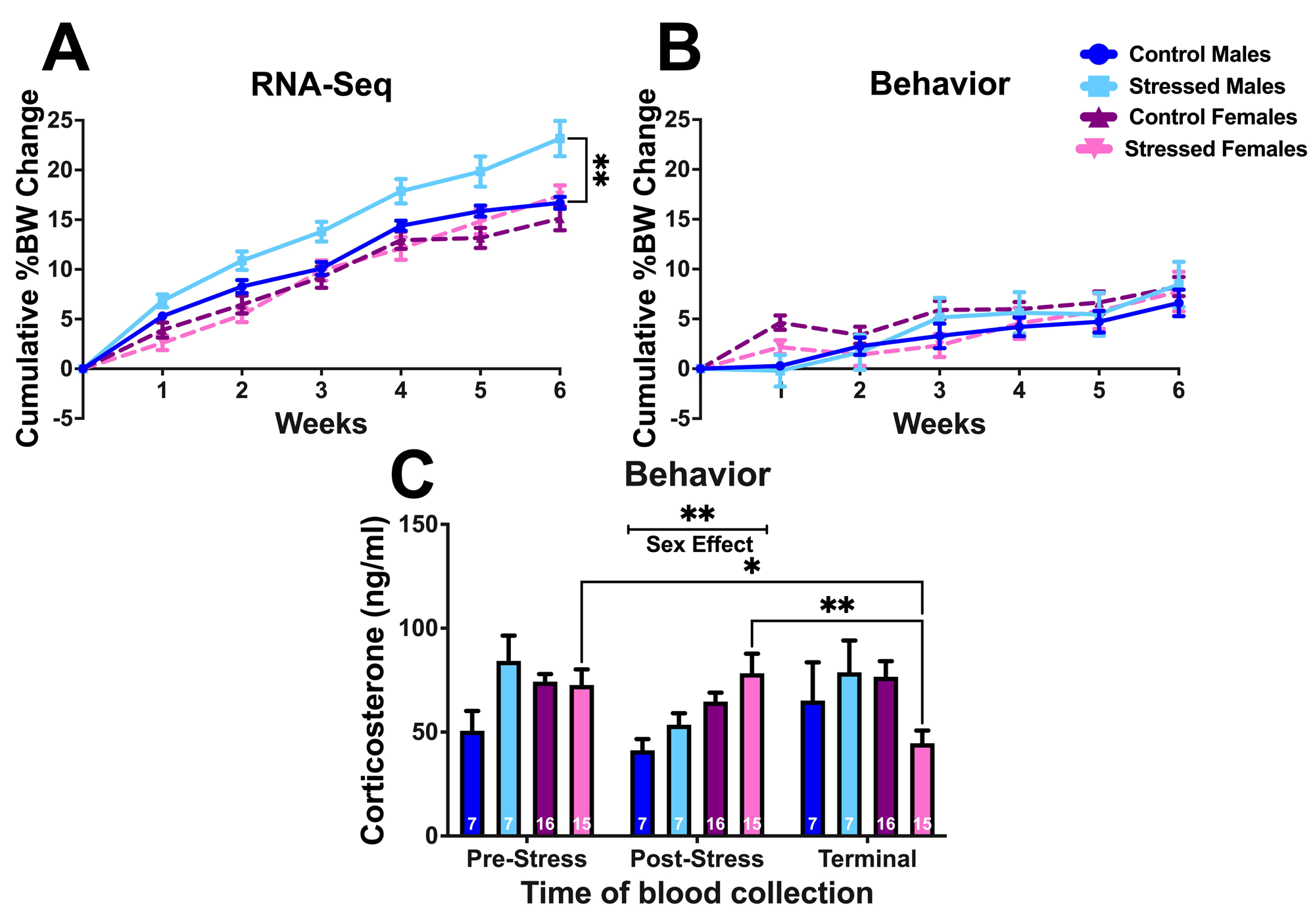
Body weight tracking of the mice used for the (A) RNA-sequencing and (B) behavior cohorts during the CSIS protocol. (C) Concentration of corticosterone in the serum of CSIS-exposed or control mice in the behavior cohorts. Data are presented as mean +/- the SEM and analyzed by a two-way ANOVA with Sidak post-hoc comparison. (* = 0.05-0.01, ** = 0.01-0.001)

For CORT assay there was no effect of stress on serum CORT concentration. For the pre-stress CORT blood, control males had an average of 50.63 ± 9.55 ng/ml, stressed males had an average of 84.28 ± 12.15 ng/ml, control females had an average of 74.28 ± 3.64 ng/ml, stressed females had an average of 72.68 ± 7.5 ng/ml (fig. 1c). There were no significant differences in treatment or sex in pre-stress serum. For post-stress, control males averaged 41.25 ± 5.37 ng/ml, stressed males averaged 53.5 ± 5.57 ng/ml, control females averaged 64.7 ± 4.29 ng/ml, and stressed females averaged 78.33 ± 9.37 ng/ml (fig. 1c). In the post-stress blood, there was no effect of treatment but there was an effect of sex in which females had significantly higher CORT levels (p = 0.0043, F (1, 34) = 9.377). For terminal blood CORT, control males averaged 65.16 ±18.42 ng/ml, stressed males averaged 78.78 ± 15.28 ng/ml, control females averaged 76.63 ± 7.5 ng/ml, and stressed females averaged 44.63 ± 6.08 ng/ml (fig. 1c). There was no effect of sex or treatment on the terminal serum concentrations. There were significant pairwise comparisons between sample collection timepoints in the females, wherein the stressed females at terminal blood collection had significantly lower concentrations of CORT than at pre-stress (p = 0.0202) and post-stress (p = 0.0037)

### 3.3 Behavior Assays

#### 3.3.1 Open Field Test

In our previous publication, the CVMS paradigm resulted in a general increase in avoidance of all stressed mice and an increase in avoidance in female mice compared to males regardless of treatment (Degroat et al., 2024). For our social stress paradigm, CSIS, we found similar but distinct effects on behavior and in relation to sex. We found no effects of stress or sex on the number of entries into the 20cm center region or the amount of time spent in the 20cm center or perimeter (fig. 2a, d, and e). In the number of entries into the perimeter, we found a main effect of treatment wherein stressed animals had fewer entries (F (1, 82) = 4.782, p = 0.0316) (fig. 2b). This suggests a reduction in exploratory behavior but does not indicate effects on avoidance. However, we did see effects on avoidance when looking at percent time spent in the corners and 10cm center. In the corners, there were pairwise effects when comparing the control males to their stressed and female counterparts as control males spent more time in the center (p = 0.0257, p = 0.0045, respectively) (fig. 2c). Interestingly, we found an interaction effect where CSIS appeared to affect avoidance behavior differently in males and females. Stress appeared to increase avoidant behaviors in males, but decreased it in females (F (1, 79) = 7.104, p = 0.0093) (fig. 2c). This interaction effect was also observed in the amount of time spent in the corners (F (1, 82) = 28.79, p < 0.0001) and is supported by pairwise comparisons that show control males spent less time in the corners compared to stressed males (p = 0.0008) but that control females spent more time in the corners compared to controls (p < 0.0001) (fig. 2f). Control males also spent more time in the center than control females (p < 0.0001) and stressed males spent less time in the center than stressed females (p = 0.0022) (fig. 2f).

**Fig. 2.**
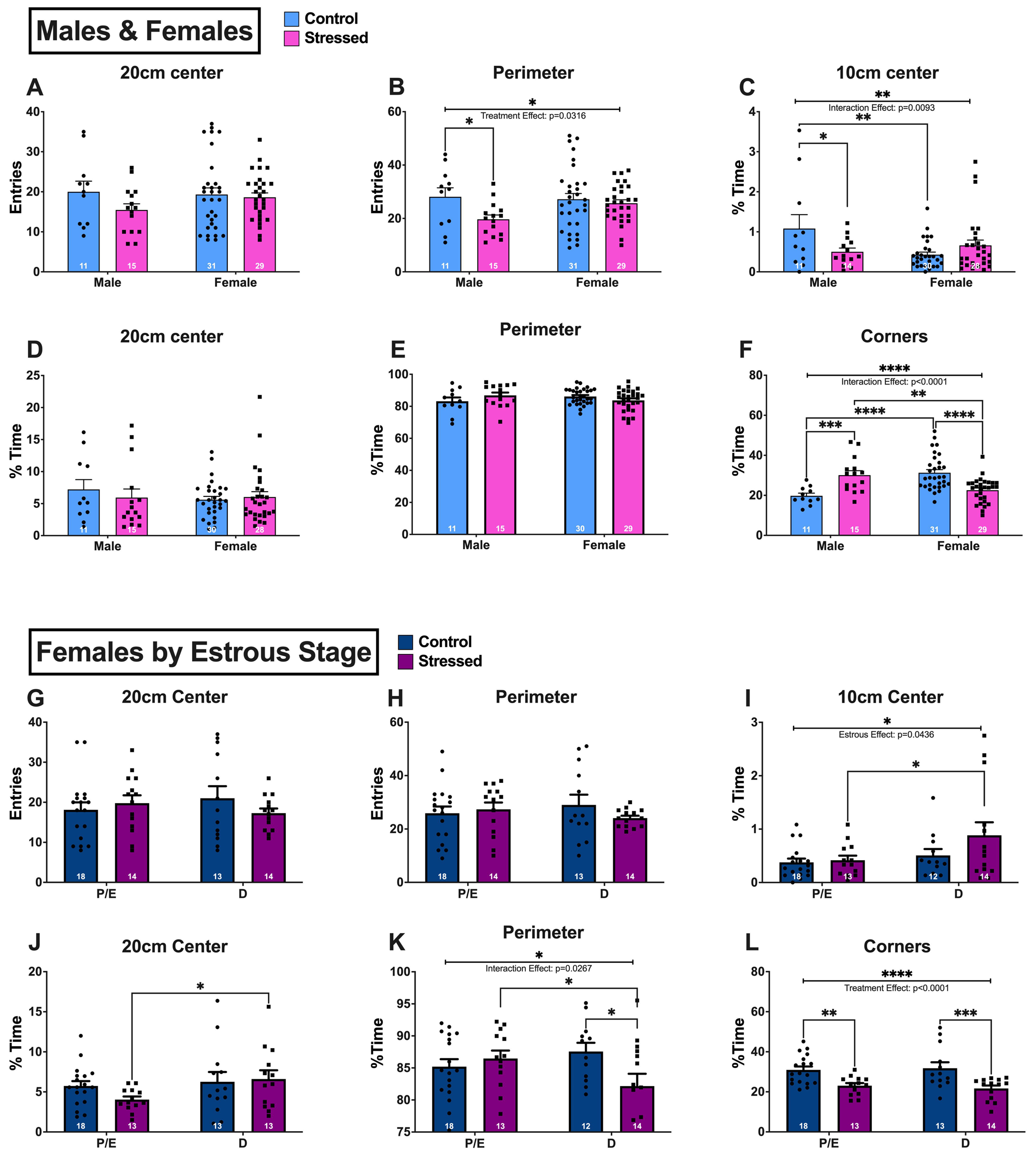
10-minute Open Field Test. (A) Distance traveled. (B) Number of entries into the 20cm center. (C) Number of entries into the perimeter. (D) Percent time spent in the 10cm center. (E) Percent time spent in the 20cm center. (F) Percent time spent in the perimeter. (G) Distance traveled with males excluded and females separated by estrus stage. (H) Number of entries into the 20cm center with males excluded and females separated by estrus stage. (I) Number of entries into the perimeter with males excluded and females separated by estrus stage. (J) Percent time spent in the 10cm center with males excluded and females separated by estrus stage. (K) Percent time spent in the 20cm center with males excluded and females separated by estrus stage. (L) Percent time spent in the perimeter with males excluded and females separated by estrus stage. Data are presented at mean +/- SEM and analyzed by two- way ANOVA with Šidák post-hoc comparisons (* = 0.05-0.01, ** = 0.01-0.001, *** = 0.001-0.0001, **** < 0.0001).

When excluding males and separating females by estrus stage, we found effects of both stress and hormone status on avoidant behaviors. We did not see significant effects in the number of entries into the 20cm or perimeter, suggesting that this type of stress does not affect exploration in the OFT (fig. 2g and h). In the amount of time spent in the 10cm center, we saw an effect of estrus stage where the D (diestrus) females spent more time in the center overall (F (1, 53) = 4.276, p = 0.0436). This is supported by a significant pairwise comparison between P/E (proetrous/estrus) stressed females and D stressed females (p = 0.0286) (fig. 2i). There was an increase in time spent in the 20cm center region in stressed D females compared to their P/E counterparts (p = 0.0478) (fig. 2j). In the perimeter, it appeared that only the diestrus females were sensitive to the stress, as we found pairwise differences in the D group such that stressed mice spent less time in the perimeter relative to controls (p = 0.0151) and D stressed females spent less time in the perimeter than P/E counterparts (p = 0.0455). This is also supported by an interaction effect (F (1, 53) = 5.195, p = 0.0267) (fig. 2k). For time spent in corner regions, we saw a main effect of stress in both groups where the stress decreased the time spent in the corners (F (1, 54) = 21.20, p < 0.0001), which is supported by pairwise comparisons between P/E control and P/E stressed mice (p = 0.0048), and D control and D stressed mice (p = 0.0008) (fig. 2l). This suggests that hormone status can affect how mice respond to stressful situations and that social stress potentially decreases avoidant behaviors in female mice.

#### 3.3.2 Elevated Plus Maze

We observed fewer effects in the EPM. We did not see any effect of sex on the number of entries into the open arms or the amount of time spent in the open arms, closed arms, or ends of the open arms (fig. 3a, d, e, and f). We saw a main effect of stress on the number of entries into the closed arms, in which stressed animals had fewer entries (F (1, 70) = 3.997, p = 0.0495) (fig. 3b). This suggests a decrease in exploratory behavior. There was also a main effect of sex in the number of entries into the ends of the open arms in which females were more willing to enter this region (F (1, 66) = 6.248, p = 0.0149) (fig. 3c). When comparing estrus stages, there were no significant differences in the entries to the closed arms or open arms ends, or amount of time spent in the open arms, closed arms or open arms ends (fig. 3h, i, j, k, and l). There was an interaction effect between stage and stress (F (1, 46) = 5.491, p = 0.0245) (fig, 3g), with stressed P/E females having fewer entries into the open arms than controls (p = 0.0246).

**Fig. 3.**
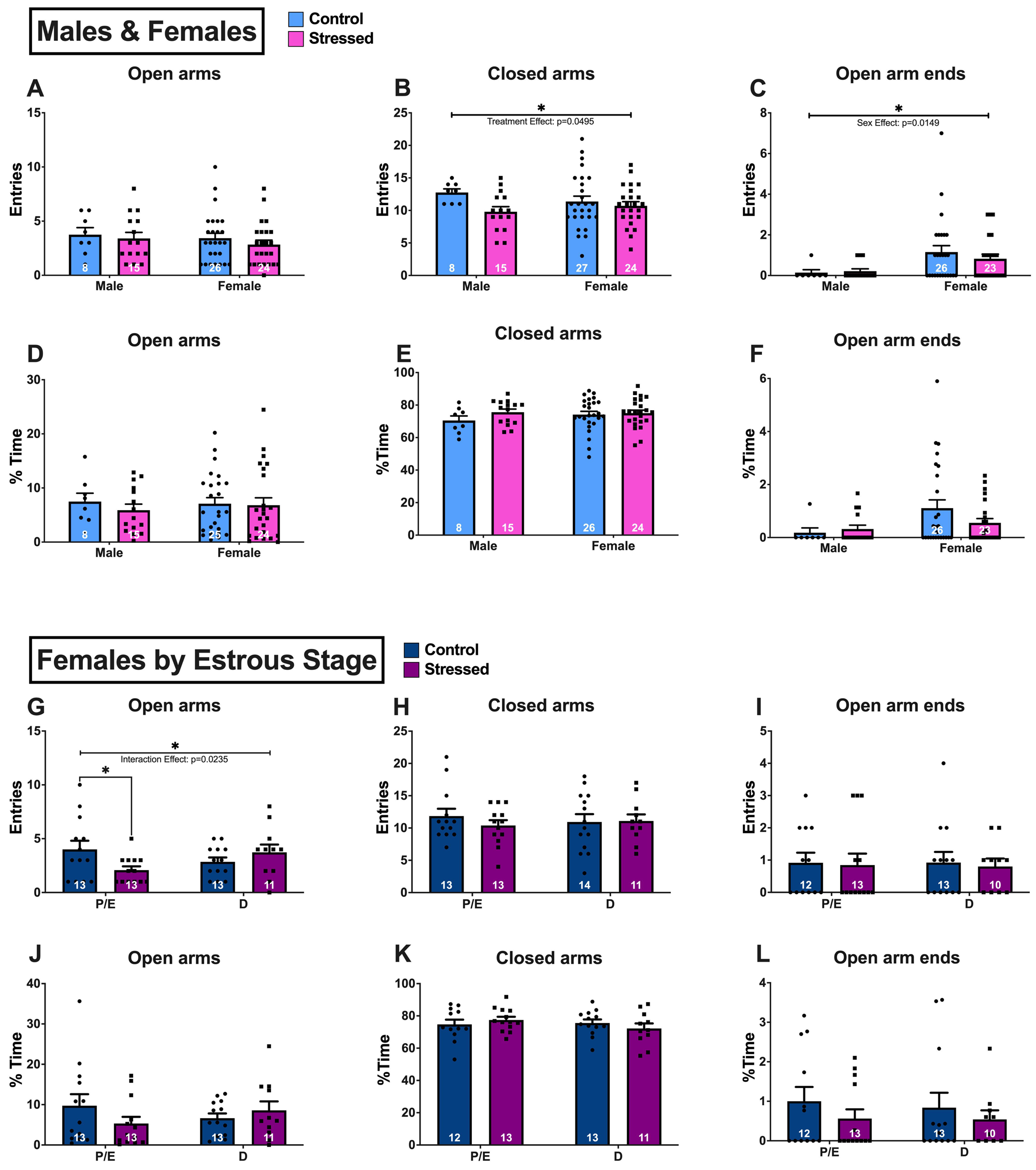
Elevated Plus Maze. (A) Distance traveled. (B) Percent time spent in open arms. (C) Number of entries into open arms. (D) Amount of time spent at the ends of the open arms. (E) Percent time spent in the closed arms. (F) Number of entries into the closed arms. (G) Distance traveled with males excluded and females separated by estrus stage. (H) Percent time spent in open arms with males excluded and females separated by estrus stage. (I) Number of entries into open arms with males excluded and females separated by estrus stage. (J) Amount of time spent at the ends of the open arms with males excluded and females separated by estrus stage. (K) Percent time spent in the closed arms with males excluded and females separated by estrus stage. (L) Number of entries into the closed arms with males excluded and females separated by estrus stage. Data are presented as mean +/- SEM and analyzed by two-way ANOVA with Šidák post-hoc comparisons (* = 0.05-0.01).

#### 3.3.3 Light/Dark Box Emergence Test

In the LDB, we saw effects in the females but not the males. We found no effects of sex or stress on the amount of time spent in the light zone (fig. 4c). We found effects of stress in the females in the number of entries into the light zone (p = 0.0456), number of entries into the transition zone (p = 0.0375), and amount of time spent in the transition zone (p = 0.0443) (fig. 4a, b, and d). Stress reduced all these measures, suggesting an increase in avoidant behaviors and a decrease in exploratory behaviors. There were no effects in the males. Within the female groups, we saw effects of both treatment and estrus stage in the LDB. In the number of entries into the light zone, we found that stressed P/E females had fewer entries compared to control P/E females (p = 0.0074) (fig. 4e). In the number of entries into the transition zone, the control P/E females had significantly more entries than both their stressed (p = 0.0056) and D (p = 0.0463) counterparts (fig. 4f). We saw a main effect of treatment on the amount of time spent in the light zone; stressed females spent less time in this zone (F (1, 54) = 6.378, p = 0.0145) (fig. 4g). There was also a pairwise stress effect between the control P/E females and their stressed counterparts (p = 0.0426) (fig. 4g). We can also see a main effect of treatment in the amount of time spent in the transition zone (F (1, 54) = 4.672, p = 0.0351) (fig. 4h). This suggests that, in contrast to the OFT, social stress increases avoidant behaviors in females as observed in the LBD.

**Fig. 4.**
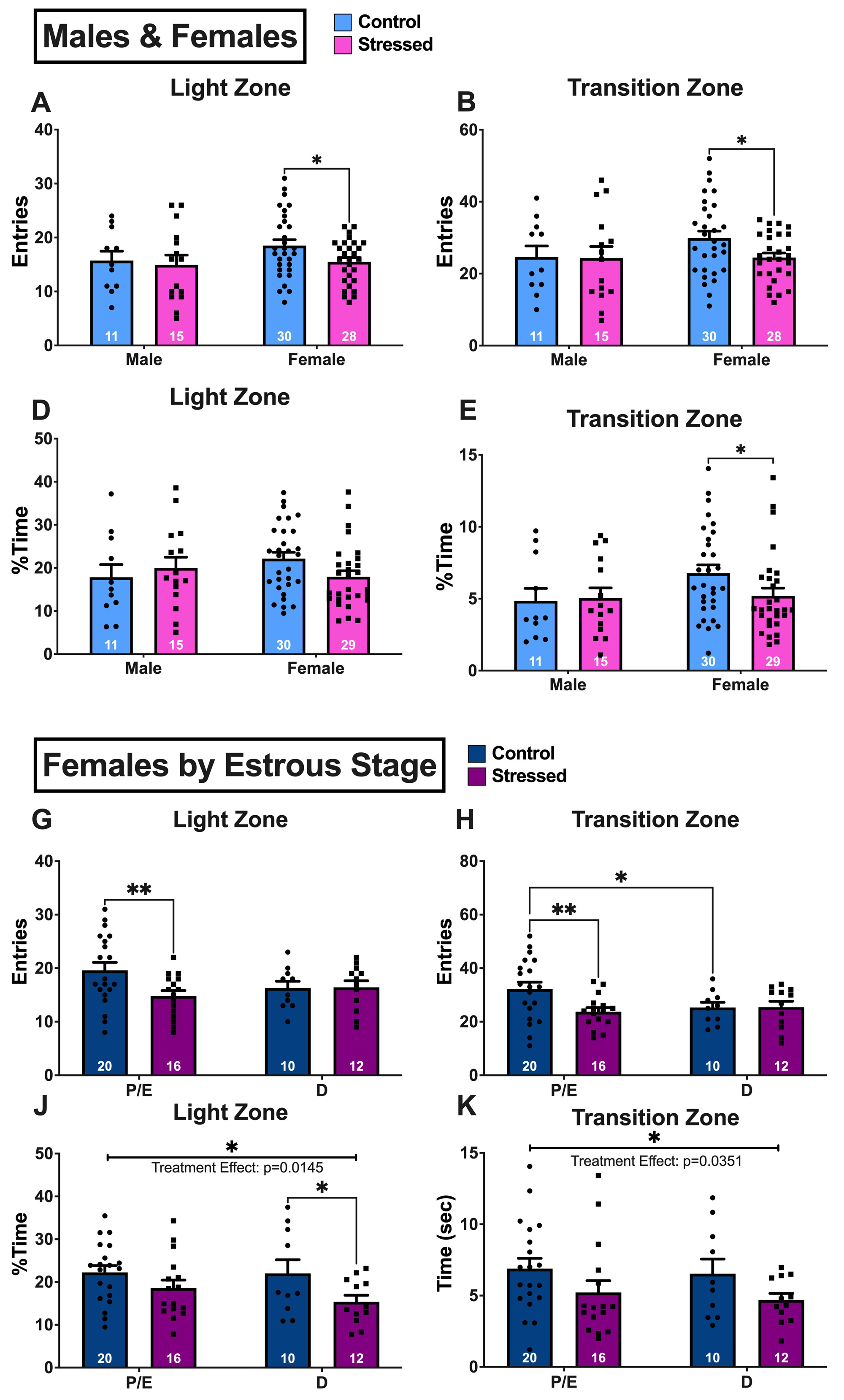
Light/Dark Box Emergence Test. (A) Distance traveled in the light and transition zones. (B) Percent time spent in the light zone. (C) Number of entries into the light zone. (D) Number of stretch attend postures while in the transition zone. (E) Percent time spent in the transition zone. (F) Number of entries into the transition zone. (G) Distance traveled in the light and transition zones with males excluded and females separated by estrus stage. (H) Percent time spent in the light zone with males excluded and females separated by estrus stage. (I) Number of entries into the light zone with males excluded and females separated by estrus stage. (J) Number of stretch attend postures while in the transition zone with males excluded and females separated by estrus stage. (K) Percent time spent in the transition zone. (L) Number of entries into the transition zone with males excluded and females separated by estrus stage. Data are presented as mean +/- SEM and analyzed by two-way ANOVA with Šidák post- hoc comparisons. (* = 0.05-0.01, ** = 0.01-0.001).

#### 3.3.4 Novelty Suppressed Feeding Test

In the NSF, we again saw both treatment and sex effects. For NSF, we tracked the amount of weight lost during the food deprivation period. We found sex effects in which the control females lost more weight than their male (p = 0.0026) and stressed (p = 0.0002) counterparts, which also resulted in a significant interaction effect between sex and stress (F (1, 83) = 11.83, p = 0.0009) (fig. 5a). In the latency to eat in the novel arena, we saw a similar interaction effect as seen in the OFT. We observed that stress caused a lower latency to eat in males (p = 0.0242) but a higher latency in females (p = 0.0175). Stressed females also had a higher latency than stressed males (p = 0.0001), resulting in a sex by stress interaction effect (F (1, 59) = 10.43, p = 0.0020) (fig. 5b). This is supported by the survival curves which show significant differences in all pairwise comparisons. Male controls were less likely to eat in the novel arena than their stressed (p = 0.0015) and female (p = 0.0459) counterparts (fig. 5c). Additionally, stressed females were less likely to eat in the novel arena than both their control (p = 0.0208) and male (p < 0.0001) counterparts (fig. 5c). This again suggests that male and female mice respond to chronic social stress in distinct ways. We did not find any effects on the amount of pellet eaten or the latency to eat in the home cage (fig. 5d and e). However, we found a main effect of sex on the probability to eat in the home cage where females were overall less likely to eat in this trial (p = 0.0088).

**Fig. 5.**
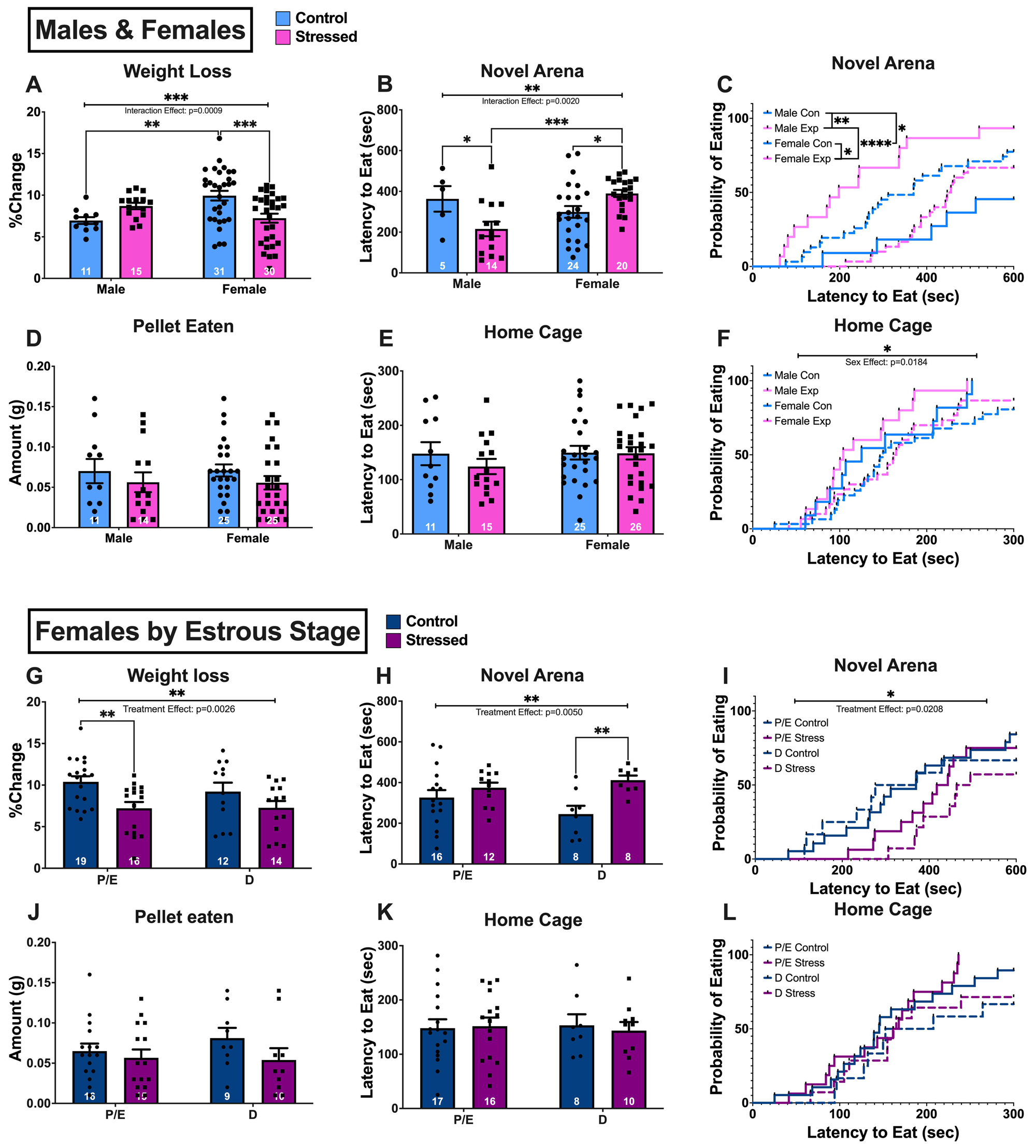
Novelty Suppressed Feeding Test. (A) Amount of weight loss during the 24-h fasting period as a percent of total body weight. (B) Latency to eat in the novel arena. (C) Kaplan-Meier survival curve for the probability to eat in the novel arena. (D) Amount of pellet eaten during test. (E) Latency to eat in the home cage. (F) Kaplan-Meier survival curve for the probability to eat in the home cage. (G) Amount of weight loss during the 24-h fasting period as a percent of total body weight with males excluded and females separated by estrus stage. (H) Latency to eat in the novel arena with males excluded and females separated by estrus stage. (I) Kaplan-Meier survival curve for the probability to eat in the novel arena with males excluded and females separated by estrus stage. (J) Amount of pellet eaten during the test with males excluded and females separated by estrus stage. (K) Latency to eat in the home cage with males excluded and females separated by estrus stage. (L) Kaplan-Meier survival curve for the probability to eat in the home cage with males excluded and females separated by estrus stage. Data are presented as mean +/- SEM. A, B, C, D, G, H, I, and J analyzed by two-way ANOVA with Šidák post-hoc comparisons. E, F, K, and L analyzed by Log-rank test (* = 0.05-0.01, ** = 0.01-0.001, *** = 0.001-0.0001, **** < 0.0001).

When excluding the males, we found effects of stress but not estrus stage. In terms of weight loss, we saw a main effect of treatment on the females where the stressed mice lost less weight than the controls (F (1, 57) = 9.902, p = 0.0026). We observed a pairwise effect of stress in the P/E females (p = 0.0041) (fig. 5g), suggesting that social stress is protective against body weight change. In the latency to eat in the novel arena, we saw a main effect of stress treatment where stressed mice had higher latencies than controls [F (1, 40) = 8.829, p = 0.0050)], along with a pairwise increase in the D females (p = 0.0062) (fig. 5h). In the survival curve for the novel arena, we saw a treatment effect where stressed animals were more likely to eat than controls (p = 0.0208) (fig. 5i). There were no effects on the amount of pellet eaten, the latency to eat in the home cage, or the probability to eat in the home cage (fig. 5j, k, and l).

### 3.4 RNA-Sequencing

In the PCA for the RNA-sequencing data, no data points fell outside the 95% confidence oval suggesting no outliers (fig. 6a). We also observed that the males and females separated out strongly. Within sexes, the male samples separated by treatment, but the females did not (fig. 6a). We found 4162 genes that were differentially expressed in the adBNST by sex (fig. 6b). 2027 genes are male-biased and 2135 are female-biased. Notable male-biased genes include: *Greb1* (log2FC = −2.0408, adj. p < 0.0001), a regulator of estrogen signaling (Mohammed et al., 2013); *Tac2* (log2FC = −0.9777, adj. p = 0.0004), neurokinin B, a peptide involved in fear learning and stress (Al Abed et al., 2021); *Mc4r* (log2FC = −0.6527, adj. p = 0.0013), melanocortin 4 receptor which responds to adrenocorticotropin and MSH hormones (Wei et al., 2023); *Htr1a* (log2FC = −0.6395, adj. p = 0.0005), a serotonin receptor (Mekli et al., 2011); *Adra2a* (log2FC = −0.5208, adj. p = 0.0005), an adrenergic receptor (Kurnik et al., 2006); *Crh* (log2FC = −0.5179, adj. p = 0.0035); *Slc1a6* (log2FC = −0.4621, adj. p = 0.0370), a glutamate transporter (Kanai and Hediger, 2004); and *Gabra3* (log2FC = 0.6295, adj. p < 0.0001), a GABA A receptor subunit implicated in depression-like behaviors in mice (Miller et al., 2010). Notable female-biased genes include: *Scn4b* (log2FC = 1.5514, adj. p < 0.0001), a subunit of voltage-gated sodium channels (Ji et al., 2017); *Drd2* (log2FC = 1.0257, adj. p = 0.0004) and *Drd1* (log2FC = 0.8513, adj. p = 0.0018), dopamine receptors (Zou et al., 2012); *Adora2a* (log2FC = 1.1594, adj. p = 0.0002), an adenosine receptor implicated in anxiety (Hohoff et al., 2020); *Camk2n1* (log2FC = 0.7756822, adj. p <0.0001) an inhibitor of CaMKII signaling pathway (Saha et al., 2007); *Syt2* (log2FC = 1.2392, adj. p < 0.0001) and *Syt6* (log2FC = 0.7613, adj. p = 0.0053), both implicated in synaptic vesicle exocytosis (Wolfes and Dean, 2020); *Inhba* (log2FC = 0.6513, p = 0.0171), a subunit of both inhibin and activin (Barton et al., 1989); *Tac1* (log2FC = 0.6117, adj. p = 0.0068), a precursor to substance P and neurokinin A (Chiwakata et al., 1991); and *Synpo* (log2FC = 0.5510, adj. p < 0.0001), a regulator of synaptic transmission (Zhang et al., 2013). When analyzing gene ontology for sex-biased differentially expressed genes, we found that male-biased genes are more likely to be related to RNA processing and protein transportation, whereas female biased genes are more likely to be related to regulation of synapses and neurogenesis (fig. 4e and f).

**Fig. 6.**
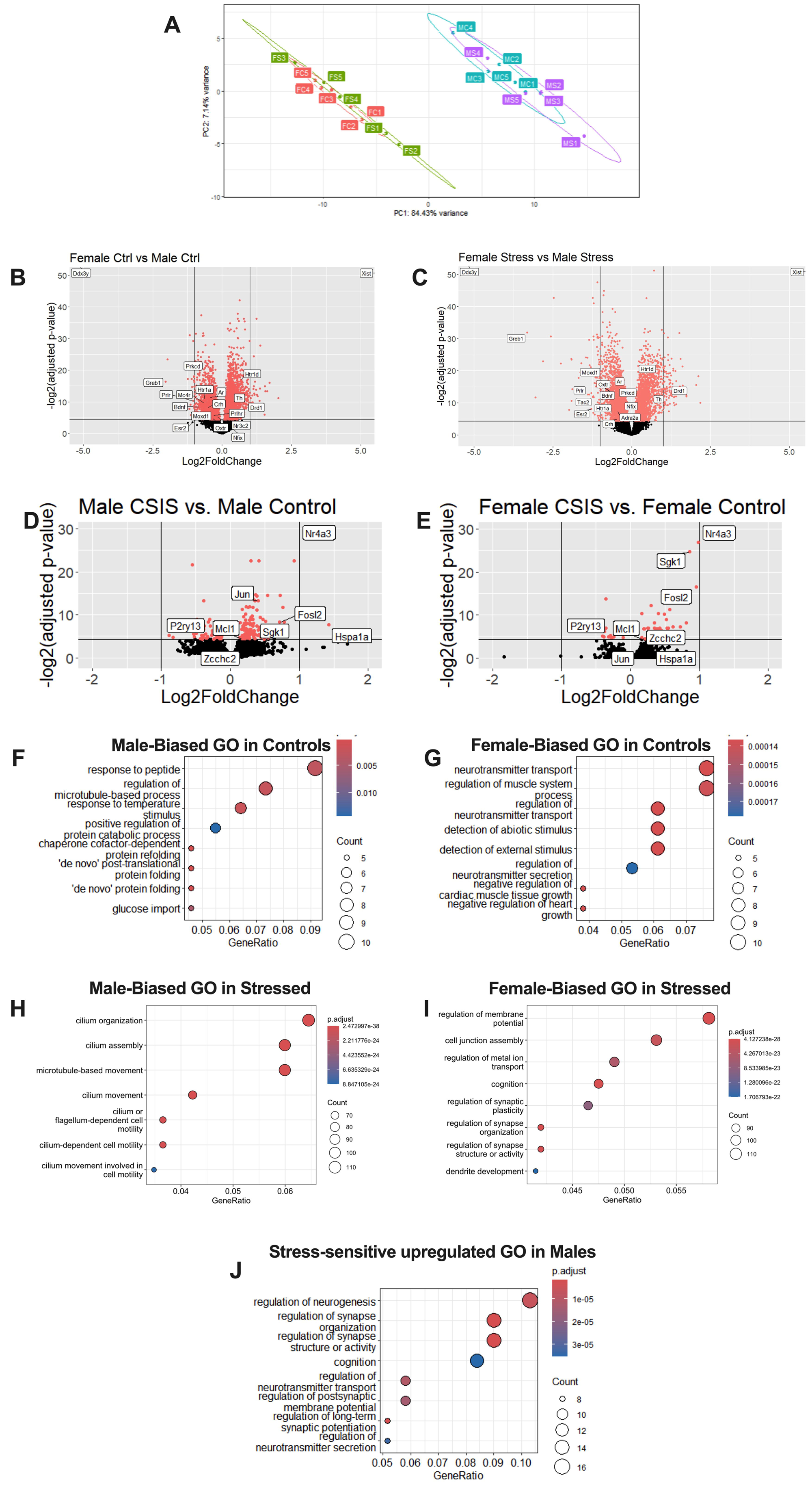
RNA-sequencing. (A) Principal component analysis (PCA), with ovals representing 95% confidence. Volcano plots comparing (B) control males to control females and (C) stressed males to stressed females. A negative fold change represents a male biased gene, and a positive fold change represents a female biased gene. (C) Volcano plot comparing control males to stressed males. A positive fold change represents an upregulation due to stress, and a negative fold change represents a downregulation due to stress. (D) Volcano plot comparing control females and stressed females. A positive fold change represents an upregulation due to stress, and a negative fold change represents a downregulation due to stress. For all volcano plots, a red dot is a significantly differentially expressed gene. (E) Gene ontology for male biased genes when comparing controls. (F) Gene ontology for female biased genes when comparing controls. (G) Gene ontology for upregulated genes due to stress in males. PCA generated with PCAExplorer, differentially expressed genes analyzed by DESeq2, Gene Ontology analyzed by clusterProfiler.

**Fig. 7.**
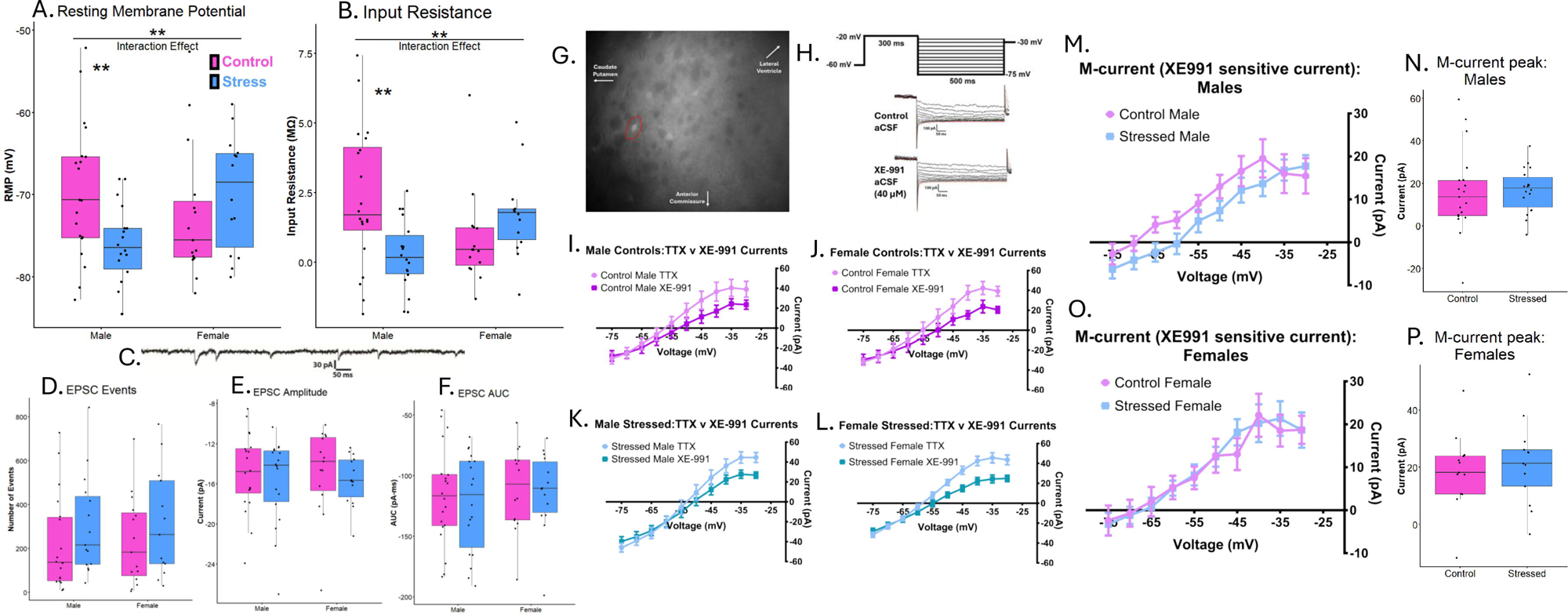
Electrophysiology of CRH+ adBNST neurons. (A) Resting membrane potential of neurons at basal conditions. (B) Input resistance of patched neurons at basal conditions. (C) Representative traces of current recordings for EPSCs. (D) Number of EPSCs detected in a 5-minute voltage-clamp recording in males and females. (E) Average amplitude of EPSCs detected in male and females. (F) Average area under the curve (AUC) of EPSCs detected in male and females. (G) Representative image of CRH+ neurons fluorescing in the adBNST. (H) M-current protocol and representative traces. (I-L) I–V plots from −75 to −30 mV before and after XE-991 perfusion in (I) control males (J) control females (K) stressed males (L) and stressed females. (M) Difference between control condition currents and currents in the presence of XE-991 – the M-current in males (N) and in females. (O) Peak M-current at −35mV in males (P) and in females. The M-current or M-current peaks were not significantly different between control and stressed groups. Boxplots (A, B, D, E, F, N, P) are depicted with median line and first and third quartile ranges in boxes, with 95 % range as whiskers. Points represent the y-axis value of each subject. significance results of Welch two-sample, two-tailed t-tests are reported (* = 0.05-0.01, ** = 0.01- 0.001). I-V plot data (I, J, K, L, M, O) are presented as mean ± SEM.

When comparing within the males, 171 genes were found to be differentially expressed due to stress (fig. 6d). 59 genes were downregulated due to stress and 112 were upregulated due to stress. Of the genes differentially expressed by stress in males, notable genes include: *Hspa1a* (log2FC = 1.4305 adj. p = 0.0049), a heat shock protein and the most highly upregulated gene (Hunt et al., 1993); *Sgk1* (log2FC = 0.4428, adj. p = 0.0395), a glucocorticoid induced kinase (Schoenebeck et al., 2005); *Synj2bp* (log2FC = 0.1675, adj. p = 0.0363), a regulator of activin signaling (Liu et al., 2006); *Mcl1* (log2FC = 0.2932, adj p < 0.0001), a regulator of apoptosis (Reed, 1998); and *Hsf2* (log2FC = 0.1974, adj. p = 0.0040), a heat shock transcription factor implicated in brain development (Chang et al., 2006). When analyzing gene ontology, genes upregulated by CSIS are more likely to be related to synaptic regulation and neuronal activity (fig. 6j). There were no gene ontologies observed to be significantly downregulated due to stress. In females, only 44 genes were found to be affected by CSIS, with 12 of those genes downregulated and 32 upregulated (fig. 6e). Notable stress sensitive genes in the females include: *Bbc3* (log2FC = −0.4071, p = 0.0318), a positive regulator of apoptosis (Han et al., 2001); *P2ry13* (log2FC = −0.3213, adj. p = 0.0324), a purinergic receptor that mediates post-synaptic acetylcholine (Guarracino et al., 2016); *Stx1b* (log2FC = −0.1026, adj. p = 0.0386), a protein involved in synaptic transmission (Mishima et al., 2021); *Hsf2* (log2FC = 0.2048, adj. p = 0.0090); and *Sgk1* (log2FC = 0.8584, adj. p < 0.0001). No significant gene ontology patterns were detected in the females. This suggests that the adBNST transcriptome in male mice may be more sensitive to social stressors.

### 3.5 Electrophysiology

#### 3.5.1 Resting state properties and M-current

In adBNST CRH neurons, there was an interaction effect between stress and sex on RMP [b = −9.350 ± 3.566, n =67, p < 0.05)] (fig. 7a). Stress significantly hyperpolarized adBNST CRH cells in males [Control: −69.6 ± 8.24; Stressed −76.2 ± 4.91; t(31.5) = 3.03, p < 0.01, d = 0.96] but had a nonsignificant depolarizing trend in females [Control: −72.5 ± 8.23; Stressed −69.7 ± 7.02]. There was also an interaction effect between sex and stress on input resistance [b = −2.9498 ± 0.9221, n =67, p < 0.01] (fig. 7b). Stress also reduced input resistance in males [Control: 2.30 ± 2.53; Stressed 0.253 ± 1.26; t(28.5) = 3.20, p < 0.01, d = 1.01] but caused a nonsignificant increase in females [Control: 0.800 ± 1.70; Stressed 1.70 ± 1.54].

As we previously characterized the M-current in oval BNST (ovBNST) CRH neurons in male mice (Hu et al., 2020), and in adBNST CRH neurons in response to chronic variable mild non-social stressors (CVMS) (Degroat et al., 2025), we next examined the M-current in CRH neurons across sexes after CSIS. XE-991 perfusion suppressed the M-current activity in control female [interaction effect (Voltage*XE-991): (9,216) = 2.880, p < 0.01] (fig. 7j), stressed female [interaction effect (Voltage*XE-991): (9,216) = 5.513, p < 0.0001] (fig. 7l), and stressed male mice [interaction effect (Voltage*XE-991): (9,306) = 3.526, p < 0.001] (fig. 7k), though it did not reach significance in control males [interaction effect (Voltage*XE-991): (9,324) = 1.376, p = 0.1977] (fig. 7i). CRH neurons from males and females did not have significant differences in the strength of their M-currents in control conditions (not shown). However, we did not find a significant interaction between stress and voltage in males [interaction effect (Voltage*XE-991): (9,315) = 0.877, p > 0.05] (fig. 7m) or females [interaction effect (Voltage*XE-991): (9,216) = 0.348, p > 0.05] (fig. 7o), nor were there effects of stress on the peak M-current in either sex (fig. 7n,p).

#### 3.5.2 EPSCs

We next examined the influence of CSIS on EPSCs in both sexes. Previous studies in our lab suggest that chronic stress augments EPSCs in male mice (Hu et al., 2020). In the current study, CSIS did not change the frequency, amplitude, or AUC of EPSCs in male or female mice. CSIS caused no difference in the number of EPSC events in males [Control: 222 ± 220; Stressed 305 ± 224] or females [Control: 234 ± 203; Stressed 314 ± 245] (fig. 7d). We also observed no difference in EPSC amplitude in males [Control: −14.9 ± 3.88 pA; Stressed −15.7 ± 4.27 pA] or females [Control: −14.8 ± 4.28 pA; Stressed −15.6 ± 2.50 pA] (fig. 7e). Likewise, there were no differences in EPSC AUC in males [Control: −117 ± 40.2; Stressed −122 ± 40.8] or females [Control: −111 ± 35.1; Stressed −113 ± 33.0] (fig. 7f).

## 4. Discussion

We sought to understand how social stress leads to neurophysiological changes that result in behavioral changes resembling mood disorders in humans. The goal of this project is to further understand the etiology of mood disorders and how it may differ between types of stressors. We first analyzed physiological markers of stress in mice, namely serum CORT concentration and body weight gain, followed by behavior assays, RNA-sequencing of the adBNST, or whole cell patch clamp electrophysiology in adBNST CRH neurons. We have shown that stress can affect mice irrespective of CORT or body weight alterations.

Initial analysis of the BW and CORT concentrations may suggest that CSIS is ineffective in females and has limited effectiveness in the males. CSIS reduced BW gain in males but not females; however, this effect was only recorded in males from one cohort. While BW is often a good correlate of stress in rodents, it is not always accurate, inhibited weight gain is seen most often in stress paradigms that use more extreme stressors (Rabasa and Dickson, 2016), nor do C57BL/J6 mice always show a BW response to chronic social stress (Moran and Delville, 2024). It is also potentially concerning that we did not observe an increase in CORT concentrations after the stress paradigms. Stress typically results in an increased basal level of CORT; however, several studies have found that CORT returns to pre-stress levels after approximately 5 weeks (Pałucha-Poniewiera et al., 2020). Our paradigm takes place over approximately 7 weeks which likely explains why we did not observe effects of stress treatment on serum CORT.

In our previous publication, we found that a chronic variable mild stress paradigm increased avoidant behaviors in general. However, in females, stress sensitivity was dependent on estrus stage, as only P/E females were sensitive to the chronic stress in part due to D females exhibiting as more avoidant regardless of stress (Degroat et al., 2024). CSIS, however, resulted in differing effects between males and females. Particularly in the OFT, social stress appeared to cause increased avoidance in the males as expected but caused the females to be less avoidant. This suggests that social stress may be protective to the females or, alternatively, reduces inhibition in females. Few studies have been conducted on the influence of stress on risk-taking behavior (Porcelli and Delgado, 2017). This is likely due the difficulty of modeling risk-taking in both humans and animals. Some studies have found that stress effects on risk-taking in humans are sex-biased, with men displaying increased risk-taking behavior after stress and women showing decreased risk-taking (Lighthall et al., 2012; Van Den Bos et al., 2009). Females’ increased OFT center time could represent an increase in risk-taking behaviors in the females, which may suggest another avenue by which to explore stress and sex. However, this stands in contrast to previous research that showed that CSIS should be similarly effective in males and females (Yohn et al., 2019). This discrepancy may be due to the different analytic approaches, as Yohn et al. analyzed distance traveled in a region in the OFT, whereas we analyzed by amount of time spent in different regions. These measures may be detecting different origins or outcomes of behavior in this assay and cannot be directly compared.

Additionally, we saw a similar sex and stress interaction in the NSF. However, for this test, the males had a shorter latency to eat in the novel arena, suggesting decreased avoidance, whereas females had a longer latency, suggesting increased avoidance. This may appear to contradict the OFT results, however, unlike the OFT, the NSF is less a test of avoidance and more a test of the interplay between avoidance and motivation, particularly food- oriented motivation. This interaction also indicates that CSIS is potentially specifically modulating motivation or risk-taking behaviors in these animals in a way that is distinct from our previous CVMS protocol.

Our behavioral test data holds interesting parallels with the electrophysiological characteristics of adBNST CRH neurons. Resting membrane potential and input resistance were significantly lower in stressed males, suggesting stress causes these cells to have a reduced likelihood to fire. As such, in males, adBNST CRH neuron activity may contribute less to ambient avoidant behaviors, while directly influencing motivated behaviors. This seems to contradict previous research in our lab showing that chemogenetic activation of anterior BNST CRH neurons reduces high effort motivated behaviors, especially in males (Maita et al., 2024a). However, this may highlight BNST CRH neurons as a distinct mediator in behaviors that require both motivation and risk-assessment, at least in males (Ch’ng et al., 2018).

Nevertheless, while the comparisons between control and stressed females for RMP and input resistance were nonsignificant, the interaction term comparing the effect of stress and sex was significant, suggesting that these neurons may play the opposite role in females. The observed changes to RMP and input resistance are also unique to social stress, as a 6-week chronic variable mild stress did not impact these metrics in male or female mice (Degroat et al., 2025).

When analyzing by estrus stage, we found that hormone status affects avoidant behaviors. Previously, we found that CVMS resulted in a very distinct effect where only P/E females appeared sensitive to stress because D females were more avoidant regardless of treatment (Degroat et al., 2024). We found more mixed results with the CSIS in the present study, where females are sensitive to some avoidant behavior elements, regardless of estrus stage, while diestrus females were less avoidant than proestrus/estrus females. This suggests that systemic and processive stress are processed and influenced differently by ovarian hormone production, wherein estrogens can both increase or decrease stressors depending on the conditions of the stress and mouse, such as P/E reducing open arm entries in the EPM in stressed but not control subjects, but D increasing OFT center time in stressed but not control subjects. This is interesting as estrogens are thought to generally decrease depressive-like symptoms in cisgender women (Payne, 2003). For example, pre-menstrual dysphoric disorder (PMDD) is a disorder very similar to major depressive disorder except that it only occurs during the transition from the follicular phase to the luteal phase of the menstrual cycle in which high estrogen levels precede a significant drop in estrogen (Hofmeister and Bodden, 2016). Recently, research in people experiencing perimenopause has found that stress sensitivity in mood is increased by fluctuations in steroid hormone concentrations and not just the absence of steroids (Eisenlohr-Moul et al., 2024; Joffe et al., 2020; Süss et al., 2021). This is a potential difference between rodents and humans and needs to be explored further.

Considering the adBNST transcriptome, we found a large difference in gene expression between the control males and control females which implies a strong sex-associated variability in the adBNST. We also saw that this sex difference was largely maintained in stressed conditions. In our gene ontology analysis, it appears that the male adBNST may be primed for transcriptional modifications, whereas the female adBNST is primed for synaptic transmission and reorganization. In males, stress seems to shift gene expression more towards motility and cilium assembly suggesting there may be differences in axonal growth and synapse formations (Jurisch-Yaksi et al., 2024). The gene function associations in males in our gene ontology analysis is interesting when considering that many more genes in the male adBNST transcriptome were impacted by stress than in females, which is similar to results we found in our transcriptomic data from a recent CVMS study (Degroat et al., 2025). In females, our gene ontology analysis suggests the adBNST is prepared for responses in synaptic transmission. On one hand, the present electrophysiological data reinforces that the male transcriptome is more sensitive to stress, as stress modifies the internal properties of adBNST CRH neurons to a hyperpolarized state, while females are less affected. Despite stress-induced changes in RMP, the M-current was not significantly altered by stress in males or females. We observed a clear lack of M-current difference in control and stressed females. In control males, XE-991 did not significantly reduce currents evoked by the step protocol, while there was a significant reduction in stressed males. This lack of XE-991 effect could partially explain an observed lack of stress-induced difference and does not necessarily rule out the M-current as a functional component of stress-induced hyperpolarization in males. Additionally, other membrane regulatory ion channels, such as the KCNK K^+^ channels which primarily maintain RMP (González et al., 2012), may also contribute to the observed changes, as some *kcnk* genes are upregulated in our stressed males. Regardless, this lack of clear M-current effect by stress is markedly different than our previously observed decrease in M-current in BNST CRH neurons in physiologically stressed males (Degroat et al., 2025). Furthermore, neither sex displayed a difference in any EPSC property, even though gene ontologies suggested that females may be more primed for synaptic reorganization. Future studies could explore inhibitory postsynaptic currents or use immunohistochemical techniques to quantify synaptic density changes within this region.

However, when looking at stress-sensitive genes, synaptic-related transcripts are significantly affected in the males, while females had far fewer differentially expressed genes (DEG). For both sexes, it may be important to understand how stress affects synaptic activity as we observed DEGs in both sexes related to synaptic transmission, such as *Synj2bp* in males or *Stx1b* in females. Previous research suggests that in the BNST, chronic stress can significantly alter synapses. Some cells, typically glutamatergic cells, to undergo long-term depression in response to chronic stress, while other cells, typically GABAergic, may increase synaptic activity (Maita et al., 2022). Additionally, apoptotic pathways should be investigated further as both males and females showed increased expression of heat-shock and apoptotic proteins, such as *Hsf2* in both sexes, *Hspa1a* in males, and *Bbc3* in females. This may also be another avenue to explore the distinct sexual differentiation of the BNST as the heat-shock proteins appear to be upregulated in stressed males and downregulated in stressed females. This could provide important insight into why males and females respond differently to stress. Additionally, it is difficult to draw distinct parallels in the present study given we used bulk RNAseq for transcriptomic analysis but focused on an individual cell population (CRH+ cells) for electrophysiology. Future RNAScope or single cell RNAseq studies could help elucidate these differences.

Overall, our study demonstrates that social stress induces different behavioral responses in male and female mice that may be due to sex variability in transcriptomic and neuronal responses to social stress in the adBNST. We have recently demonstrated that chemogenetic activation of CRH neurons in the adBNST reduces motivation and avoidance behaviors with subtle variability across sexes (Maita et al., 2024b). However, these results require further exploration as females appeared more resistant to the social stressors in terms of effects on avoidance behaviors. Future studies will characterize the influence of CSIS on motivational behaviors controlled by adBNST CRH neurons, especially those involving a risk-assessment component. Future studies should further characterize the transcriptional and molecular mechanisms underlying sex variability in the adBNST. In summary, we have demonstrated pathways through which stress is differentially regulated between male and female adBNST, which may have translational implications for sex differences in diagnoses and prognoses of stress-related mood disorders.

## Acknowledgements

This research is funded by NIH/NIMH R01MH123544 (T.A.R. and B.S) and 5T32ES007148 (K.M.)

## Author Contributions

**Thomas J Degroat*:** Formal Analysis, Investigation, Data Curation, Writing - Original Draft, Project Administration. **Kevin M Moran*:** Formal Analysis, Investigation, Data Curation, Writing - Original Draft. **Katherine Denney:** Formal Analysis **Sarah Paladino:** Data Curation. **Ali Yasrebi:** Investigation. **Nadja Knox:** Investigation. **Benjamin A Samuels:** Conceptualization, Methodology, Writing - Review & Editing, Funding Acquisition. **Troy A Roepke:** Conceptualization, Methodology, Resources, Writing - Review & Editing, Supervision, Project Administration, Funding Acquisition.

## Data Availability Statement

The data that support the findings of this study are available on request from the corresponding author.

---

**Table 1.**
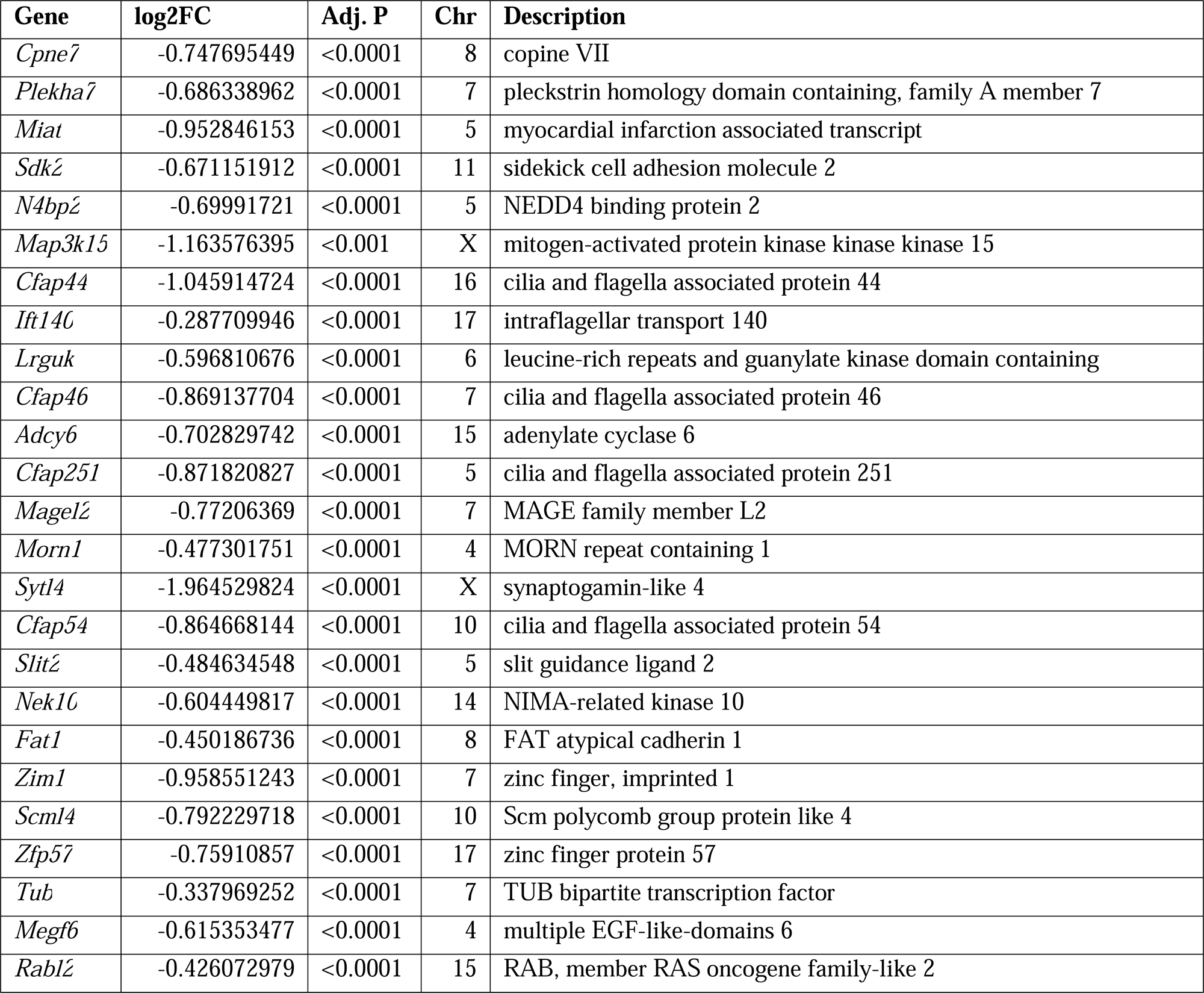
Top 25 most significant male-biased genes in control groups that are not Y-linked. Genes are listed in order of the lowest adjusted p-value alongside log2 fold change (FC), chromosome location (chr) and description of the gene. A positive log2FC represents an upregulation due to stress. A negative log_2_ fold change represents a downregulation due to stress.

**Table 2.**
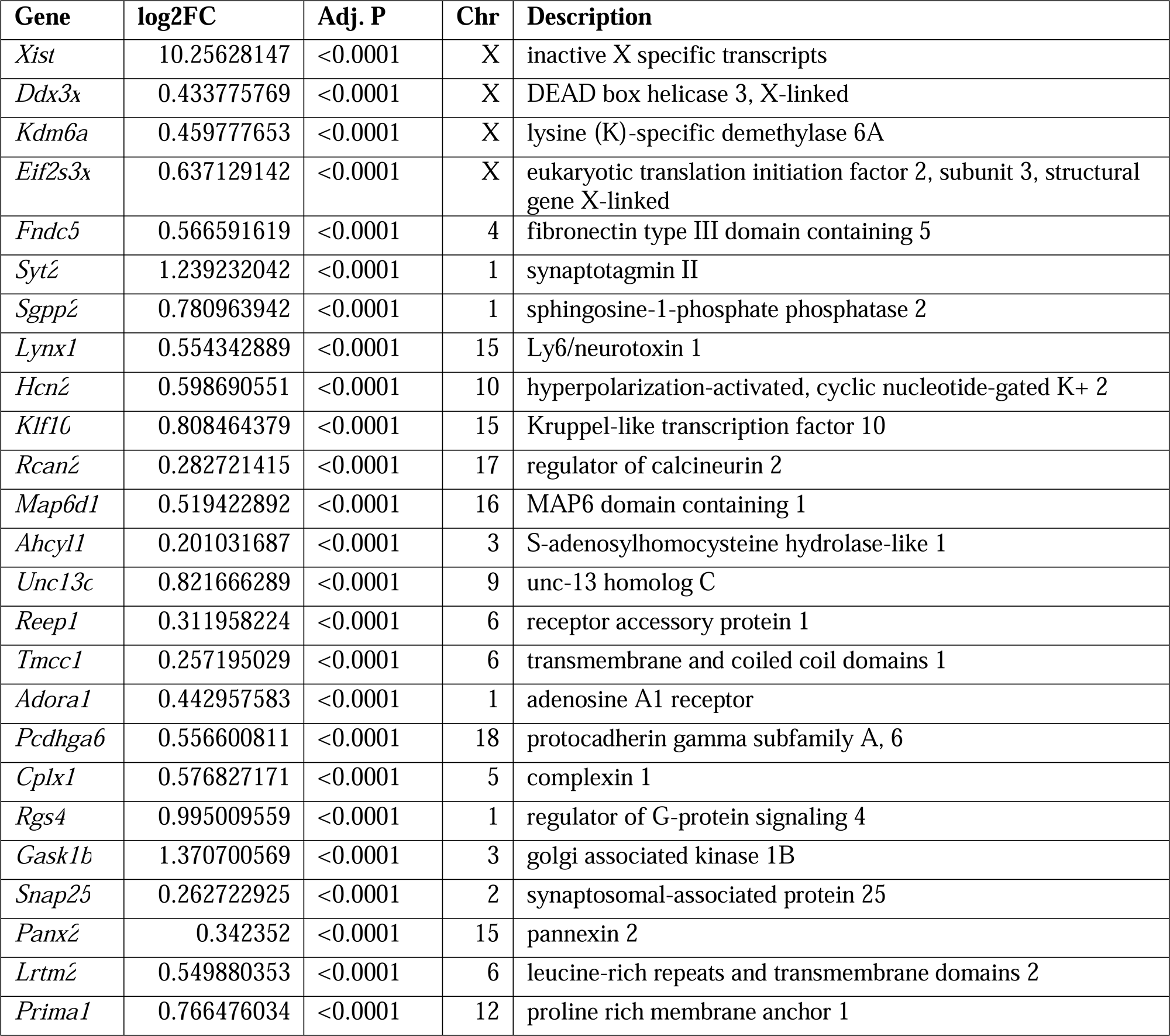
Top 25 most significant female-biased genes in control groups. Genes are listed in order of the lowest adjusted p-value alongside log2 fold change (FC), chromosome location (chr) and description of the gene. A positive log2FC represents an upregulation due to stress. A negative log_2_ fold change represents a downregulation due to stress.

**Table 3.**
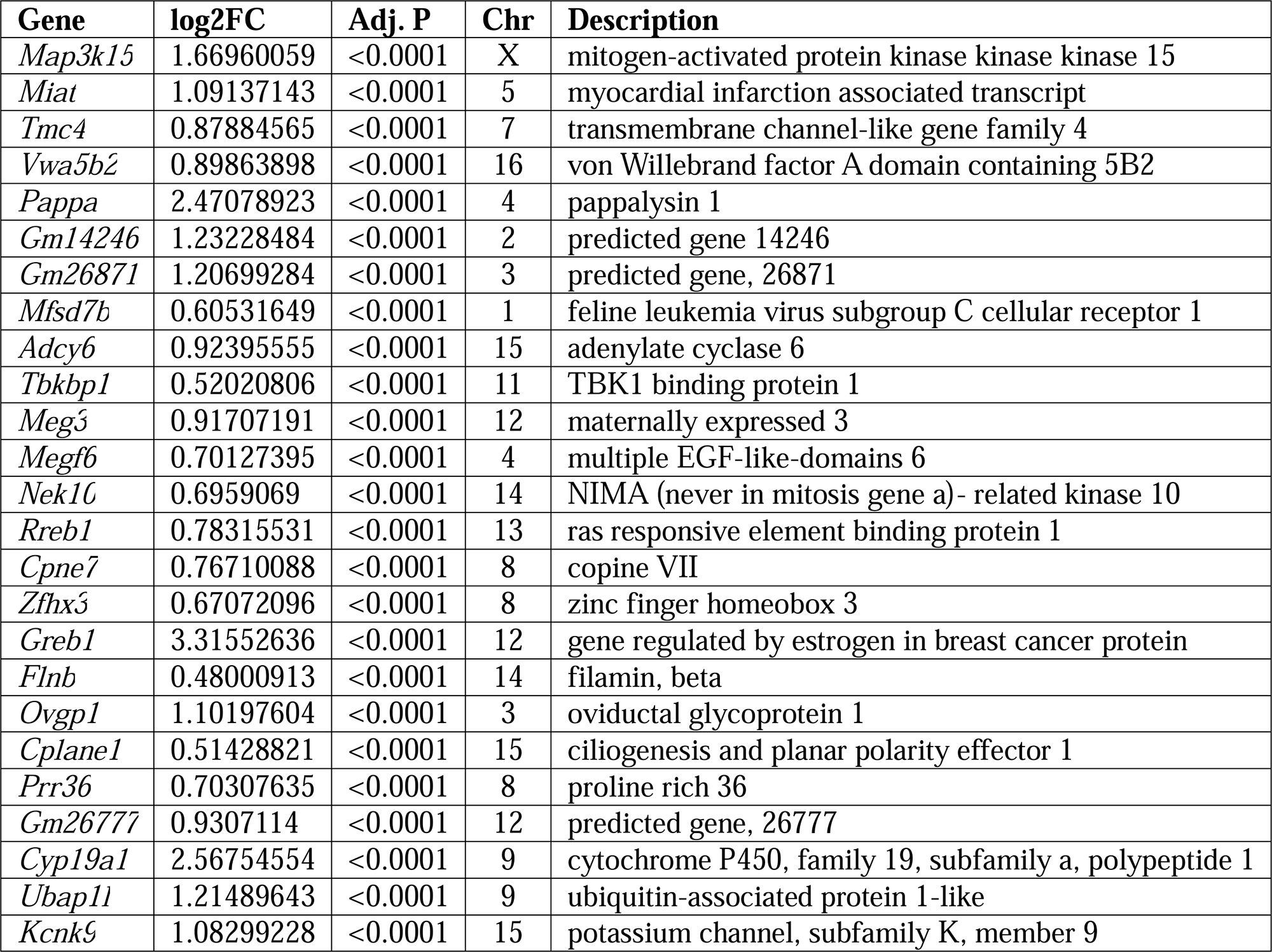
Top 25 most significant male-biased genes in stressed groups that are not Y-linked. Genes are listed in order of the lowest adjusted p-value alongside log2 fold change (FC), chromosome location (chr) and description of the gene. A positive log2FC represents an upregulation due to stress. A negative log_2_ fold change represents a downregulation due to stress.

**Table 4.**
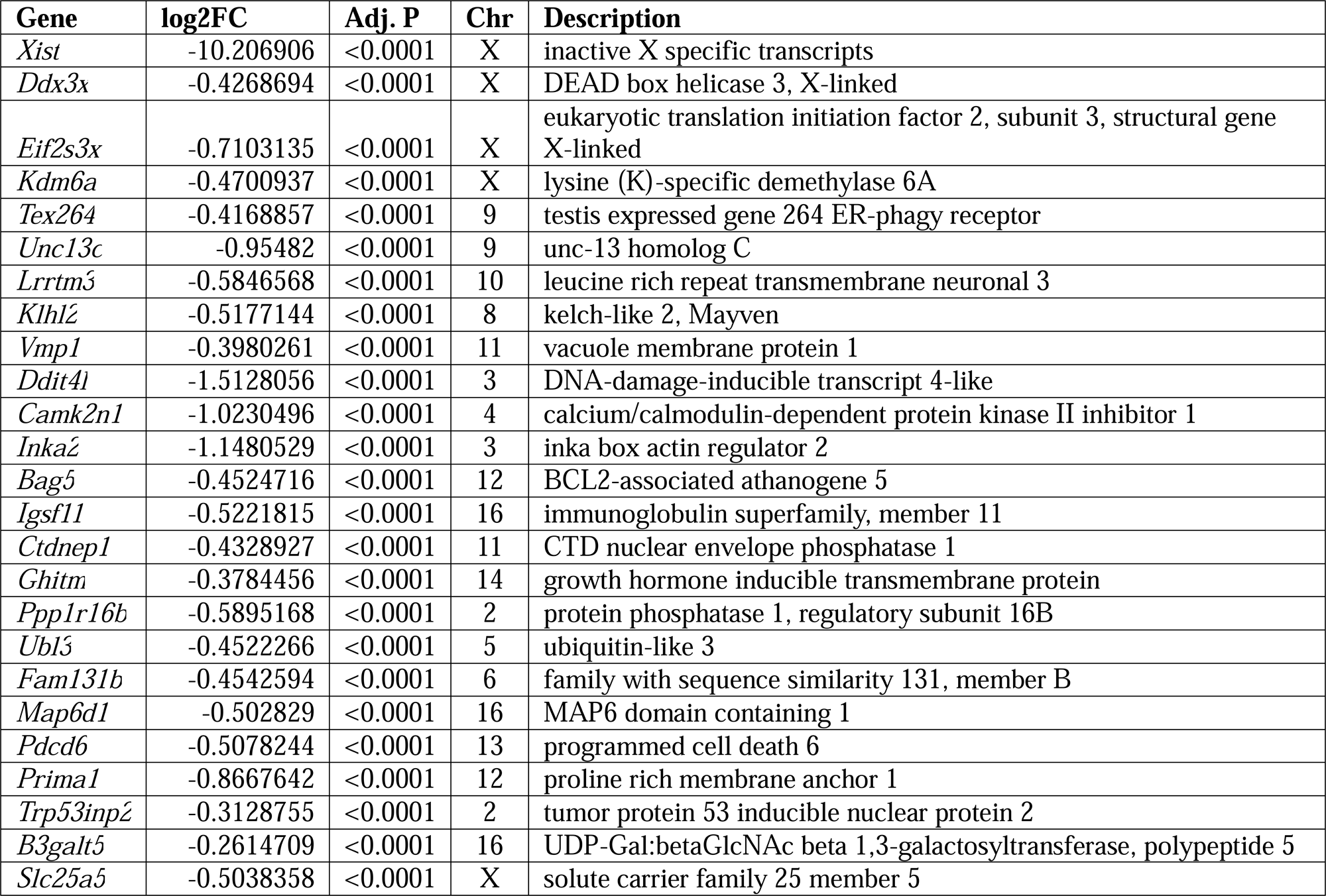
Top 25 most significant female-biased genes in stressed groups. Genes are listed in order of the lowest adjusted p-value alongside log2 fold change (FC), chromosome location (chr) and description of the gene. A positive log2FC represents an upregulation due to stress. A negative log_2_ fold change represents a downregulation due to stress.

**Table 5.**
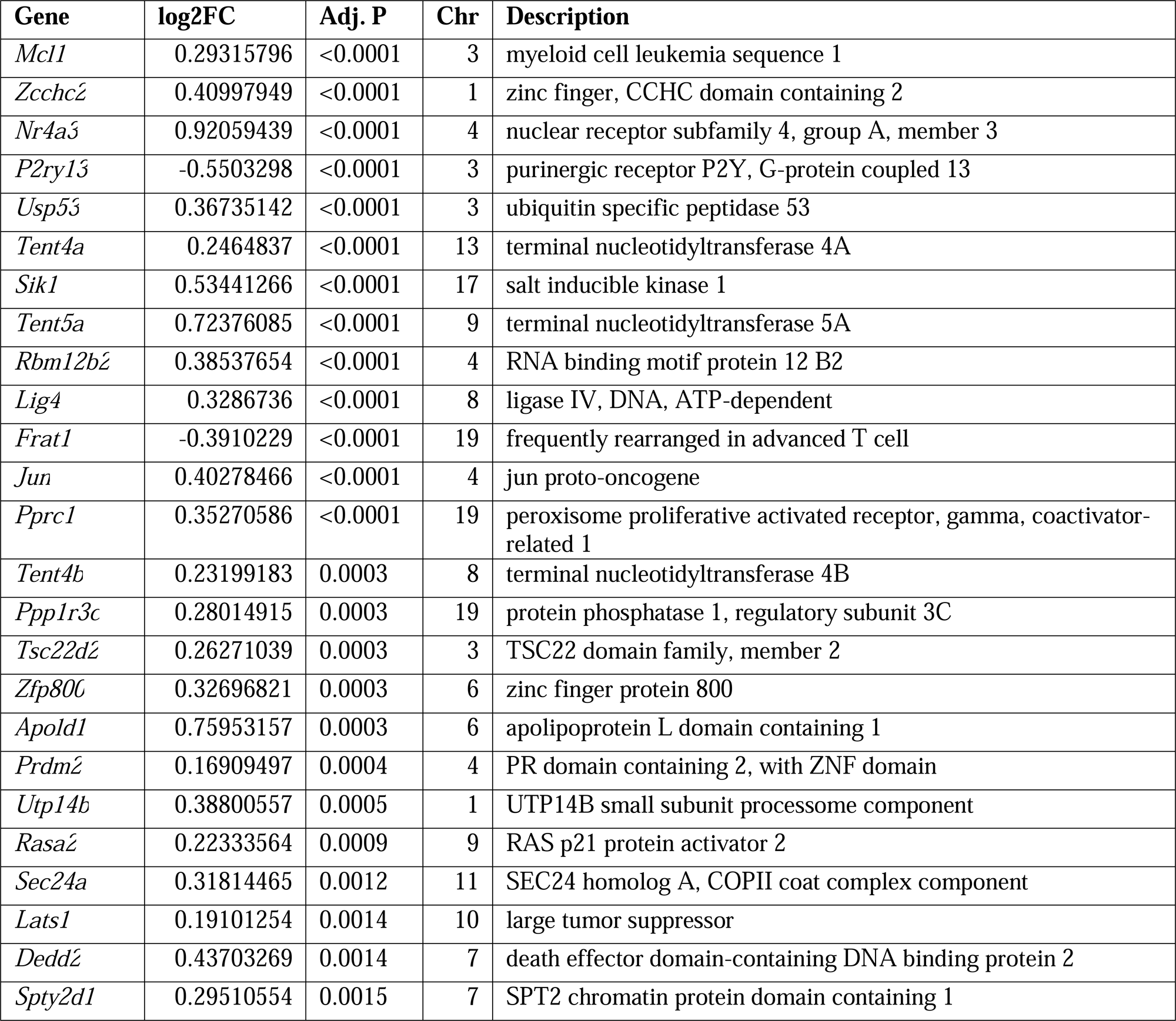
Top 25 most significant stress-sensitive genes in males ordered by adjusted p-value. Genes are listed in order of the lowest adjusted p-value alongside log2 fold change (FC), chromosome location (chr) and description of the gene. A positive log2FC represents an upregulation due to stress. A negative log_2_ fold change represents a downregulation due to stress.

**Table 6.**
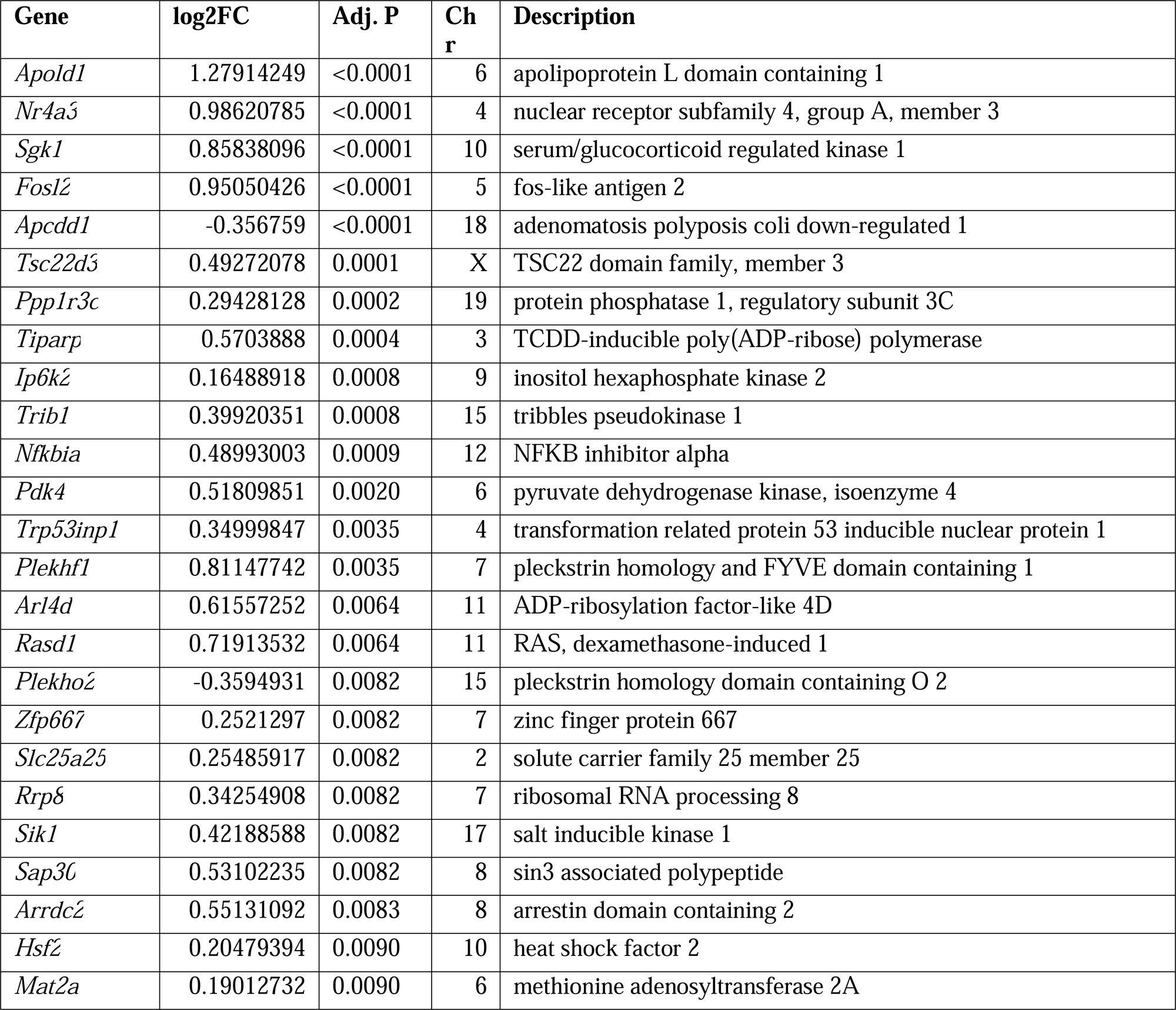
Top 25 most significant stress-sensitive genes in females. Genes are listed in order of the lowest adjusted p- value alongside log2 fold change (FC), chromosome location (chr) and description of the gene. A positive log2FC represents an upregulation due to stress. A negative log_2_ fold change represents a downregulation due to stress.

## References

Al Abed, A.S., Reynolds, N.J., Dehorter, N., 2021. A Second Wave for the Neurokinin Tac2 Pathway in Brain Research. Biol. Psychiatry, BPS 90/3 Persisting Neural Impact of Social Adversity 90, 156–164. 10.1016/j.biopsych.2021.02.016

Allen, L.S., Gorski, R.A., 1990. Sex difference in the bed nucleus of the stria terminalis of the human brain. J. Comp. Neurol. 302, 697–706. 10.1002/cne.903020402

Barton, D.E., Yang-Feng, T.L., Mason, A.J., Seeburg, P.H., Francke, U., 1989. Mapping of genes for inhibin subunits α, βA, and βB on human and mouse chromosomes and studies of *jsd* mice. Genomics 5, 91–99. 10.1016/0888-7543(89)90091-8

Beyeler, A., Dabrowska, J., 2020. Neuronal diversity of the amygdala and the bed nucleus of the stria terminalis. Handb. Behav. Neurosci. 26, 63–100. 10.1016/b978-0-12-815134-1.00003-9

Brown, D.A., Adams, P.R., 1980. Muscarinic suppression of a novel voltage-sensitive K+ current in a vertebrate neurone. Nature 283, 673–676. 10.1038/283673a0

Chang, Y., Östling, P., Åkerfelt, M., Trouillet, D., Rallu, M., Gitton, Y., El Fatimy, R., Fardeau, V., Le Crom, S., Morange, M., Sistonen, L., Mezger, V., 2006. Role of heat-shock factor 2 in cerebral cortex formation and as a regulatorof p35 expression. Genes Dev. 20, 836–847. 10.1101/gad.366906

Chiwakata, C., Brackmann, B., Hunt, N., Davidoff, M., Schulze, W., Ivell, R., 1991. Tachykinin (Substance-P) Gene Expression in Leydig Cells of the Human and Mouse Testis. Endocrinology 128, 2441–2448. 10.1210/endo-128-5-2441

Ch’ng, S., Fu, J., Brown, R.M., McDougall, S.J., Lawrence, A.J., 2018. The intersection of stress and reward: BNST modulation of aversive and appetitive states. Prog. Neuropsychopharmacol. Biol. Psychiatry, New directions in addiction 87, 108–125. 10.1016/j.pnpbp.2018.01.005

Degroat, T.J., Paladino, S.E., Samuels, B.A., Roepke, T.A., 2025. Chronic variable mild stress alters the transcriptome and signaling properties of the anterodorsal bed nucleus of the stria terminalis in a sex-dependent manner. J. Neuroendocrinol. 37, e70041. 10.1111/jne.70041

Degroat, T.J., Wiersielis, K., Denney, K., Kodali, S., Daisey, S., Tollkuhn, J., Samuels, B.A., Roepke, T.A., 2024. Chronic stress and its effects on behavior, RNA expression of the bed nucleus of the stria terminalis, and the M-current of NPY neurons. Psychoneuroendocrinology 161, 106920. 10.1016/j.psyneuen.2023.106920

Dobin, A., Davis, C.A., Schlesinger, F., Drenkow, J., Zaleski, C., Jha, S., Batut, P., Chaisson, M., Gingeras, T.R., 2013. STAR: ultrafast universal RNA-seq aligner. Bioinformatics 29, 15–21. 10.1093/bioinformatics/bts635

Eisenlohr-Moul, T., Swales, D.A., Rubinow, D.R., Schiff, L., Schiller, C.E., 2024. Temporal dynamics of neurobehavioral hormone sensitivity in a scaled-down experimental model of early pregnancy and parturition. Neuropsychopharmacology 49, 414–421. 10.1038/s41386-023-01687-0

Flanigan, M.E., Kash, T.L., 2022. Coordination of social behaviors by the bed nucleus of the stria terminalis. Eur. J. Neurosci. 55, 2404–2420. 10.1111/ejn.14991

Gegenhuber, B., Wu, M.V., Bronstein, R., Tollkuhn, J., 2022. Gene regulation by gonadal hormone receptors underlies brain sex differences. Nature 606, 153–159. 10.1038/s41586-022-04686-1

Golden, S.A., Covington, H.E., Berton, O., Russo, S.J., 2011. A standardized protocol for repeated social defeat stress in mice. Nat. Protoc. 6, 1183–1191. 10.1038/nprot.2011.361

González, C., Baez-Nieto, D., Valencia, I., Oyarzún, I., Rojas, P., Naranjo, D., Latorre, R., 2012. K+ Channels: Function- Structural Overview. Compr. Physiol. 2, 2087–2149. 10.1002/j.2040-4603.2012.tb00452.x

Guarracino, J.F., Cinalli, A.R., Fernández, V., Roquel, L.I., Losavio, A.S., 2016. P2Y13 receptors mediate presynaptic inhibition of acetylcholine release induced by adenine nucleotides at the mouse neuromuscular junction. Neuroscience 326, 31–44. 10.1016/j.neuroscience.2016.03.066

Han, J., Flemington, C., Houghton, A.B., Gu, Z., Zambetti, G.P., Lutz, R.J., Zhu, L., Chittenden, T., 2001. Expression of *bbc3*, a pro-apoptotic BH3-only gene, is regulated by diverse cell death and survival signals. Proc. Natl. Acad. Sci. 98, 11318–11323. 10.1073/pnas.201208798

Harris, A.Z., Atsak, P., Bretton, Z.H., Holt, E.S., Alam, R., Morton, M.P., Abbas, A.I., Leonardo, E.D., Bolkan, S.S., Hen, R., Gordon, J.A., 2018. A Novel Method for Chronic Social Defeat Stress in Female Mice. Neuropsychopharmacology 43, 1276–1283. 10.1038/npp.2017.259

Herman, J.P., Cullinan, W.E., 1997. Neurocircuitry of stress: central control of the hypothalamo–pituitary–adrenocortical axis. Trends Neurosci. 20, 78–84. 10.1016/S0166-2236(96)10069-2

Hofmeister, S., Bodden, S., 2016. Premenstrual Syndrome and Premenstrual Dysphoric Disorder. Am. Fam. Physician 94, 236–240.

Hohoff, C., Kroll, T., Zhao, B., Kerkenberg, N., Lang, I., Schwarte, K., Elmenhorst, D., Elmenhorst, E.-M., Aeschbach, D., Zhang, W., Baune, B.T., Neumaier, B., Bauer, A., Deckert, J., 2020. ADORA2A variation and adenosine A1 receptor availability in the human brain with a focus on anxiety-related brain regions: modulation by ADORA1 variation. Transl. Psychiatry 10, 406. 10.1038/s41398-020-01085-w

Hu, P., Maita, I., Phan, M.L., Gu, E., Kwok, C., Dieterich, A., Gergues, M.M., Yohn, C.N., Wang, Y., Zhou, J.-N., Qi, X.-R., Swaab, D.F., Pang, Z.P., Lucassen, P.J., Roepke, T.A., Samuels, B.A., 2020. Early-life stress alters affective behaviors in adult mice through persistent activation of CRH-BDNF signaling in the oval bed nucleus of the stria terminalis. Transl. Psychiatry 10, 396. 10.1038/s41398-020-01070-3

Hunt, C.R., Gasser, D.L., Chaplin, D.D., Pierce, J.C., Kozak, C.A., 1993. Chromosomal Localization of Five Murine HSP70 Gene Family Members: Hsp70.1, Hsp70-2, Hsp70-3, Hsc70t, and Grp78. Genomics 16, 193–198. 10.1006/geno.1993.1158

Ji, X., Saha, S., Gao, G., Lasek, A.W., Homanics, G.E., Guildford, M., Tapper, A.R., Martin, G.E., 2017. The Sodium Channel β4 Auxiliary Subunit Selectively Controls Long-Term Depression in Core Nucleus Accumbens Medium Spiny Neurons. Front. Cell. Neurosci. 11. 10.3389/fncel.2017.00017

Joffe, H., de Wit, A., Coborn, J., Crawford, S., Freeman, M., Wiley, A., Athappilly, G., Kim, S., Sullivan, K.A., Cohen, L.S., Hall, J.E., 2020. Impact of Estradiol Variability and Progesterone on Mood in Perimenopausal Women With Depressive Symptoms. J. Clin. Endocrinol. Metab. 105, e642–e650. 10.1210/clinem/dgz181

Jurisch-Yaksi, N., Wachten, D., Gopalakrishnan, J., 2024. The neuronal cilium – a highly diverse and dynamic organelle involved in sensory detection and neuromodulation. Trends Neurosci. 47, 383–394. 10.1016/j.tins.2024.03.004

Kanai, Y., Hediger, M.A., 2004. The glutamate/neutral amino acid transporter family SLC1: molecular, physiological and pharmacological aspects. Pflüg. Arch. 447, 469–479. 10.1007/s00424-003-1146-4

Kinn Rød, A.M., Milde, A.M., Grønli, J., Jellestad, F.K., Sundberg, H., Murison, R., 2012. Long-term effects of footshock and social defeat on anxiety-like behaviours in rats: Relationships to pre-stressor plasma corticosterone concentration. Stress 15, 658–670. 10.3109/10253890.2012.663836

Kogler, L., Müller, V.I., Chang, A., Eickhoff, S.B., Fox, P.T., Gur, R.C., Derntl, B., 2015. Psychosocial versus physiological stress — Meta-analyses on deactivations and activations of the neural correlates of stress reactions. NeuroImage 119, 235–251. 10.1016/j.neuroimage.2015.06.059

Krahn, D.D., Gosnell, B.A., Majchrzak, M.J., 1990. The anorectic effects of CRH and restraint stress decrease with repeated exposures. Biol. Psychiatry 27, 1094–1102. 10.1016/0006-3223(90)90046-5

Kurnik, D., Muszkat, M., Li, C., Sofowora, G.G., Solus, J., Xie, H.-G., Harris, P.A., Jiang, L., McMunn, C., Ihrie, P., Dawson, E.P., Williams, S.M., Wood, A.J.J., Stein, C.M., 2006. Variations in the α2A-adrenergic receptor gene and their functional effects. Clin. Pharmacol. Ther. 79, 173–185. 10.1016/j.clpt.2005.10.006

Liao, Y., Smyth, G.K., Shi, W., 2014. featureCounts: an efficient general purpose program for assigning sequence reads to genomic features. Bioinformatics 30, 923–930. 10.1093/bioinformatics/btt656

Lighthall, N.R., Sakaki, M., Vasunilashorn, S., Nga, L., Somayajula, S., Chen, E.Y., Samii, N., Mather, M., 2012. Gender differences in reward-related decision processing under stress. Soc. Cogn. Affect. Neurosci. 7, 476–484. 10.1093/scan/nsr026

Liu, Z.H., Tsuchida, K., Matsuzaki, T., Bao, Y.L., Kurisaki, A., Sugino, H., 2006. Characterization of isoforms of activin receptor-interacting protein 2 that augment activin signaling. J. Endocrinol. 189, 409–421. 10.1677/joe.1.06420

Love, M.I., Huber, W., Anders, S., 2014. Moderated estimation of fold change and dispersion for RNA-seq data with DESeq2. Genome Biol. 15, 550. 10.1186/s13059-014-0550-8

Maita, I., Bazer, A., Chae, K., Parida, A., Mirza, M., Sucher, J., Phan, M., Liu, T., Hu, P., Soni, R., Roepke, T.A., Samuels, B.A., 2024a. Chemogenetic activation of corticotropin-releasing factor-expressing neurons in the anterior bed nucleus of the stria terminalis reduces effortful motivation behaviors. Neuropsychopharmacology 49, 377–385. 10.1038/s41386-023-01646-9

Maita, I., Bazer, A., Chae, K., Parida, A., Mirza, M., Sucher, J., Phan, M., Liu, T., Hu, P., Soni, R., Roepke, T.A., Samuels, B.A., 2024b. Chemogenetic activation of corticotropin-releasing factor-expressing neurons in the anterior bed nucleus of the stria terminalis reduces effortful motivation behaviors. Neuropsychopharmacology 49, 377–385. 10.1038/s41386-023-01646-9

Maita, I., Roepke, T.A., Samuels, B.A., 2022. Chronic stress-induced synaptic changes to corticotropin-releasing factor- signaling in the bed nucleus of the stria terminalis. Front. Behav. Neurosci. 16.

Marini, F., Binder, H., 2019. pcaExplorer: an R/Bioconductor package for interacting with RNA-seq principal components. BMC Bioinformatics 20, 331. 10.1186/s12859-019-2879-1

Marrion, N.V., 1997. Control of M-current. Annu. Rev. Physiol. 59, 483–504. 10.1146/annurev.physiol.59.1.483

Mekli, K., Payton, A., Miyajima, F., Platt, H., Thomas, E., Downey, D., Lloyd-Williams, K., Chase, D., Toth, Z.G., Elliott, R., Ollier, W.E., Anderson, I.M., Deakin, J.F.W., Bagdy, G., Juhasz, G., 2011. The HTR1A and HTR1B receptor genes influence stress-related information processing. Eur. Neuropsychopharmacol. 21, 129–139. 10.1016/j.euroneuro.2010.06.013

Miller, B.H., Schultz, L.E., Long, B.C., Pletcher, M.T., 2010. Quantitative trait locus analysis identifies Gabra3 as a regulator of behavioral despair in mice. Mamm. Genome 21, 247–257. 10.1007/s00335-010-9266-6

Mishima, T., Fujiwara, T., Kofuji, T., Saito, A., Terao, Y., Akagawa, K., 2021. Syntaxin 1B regulates synaptic GABA release and extracellular GABA concentration, and is associated with temperature-dependent seizures. J. Neurochem. 156, 604–613. 10.1111/jnc.15159

Mohammed, H., D’Santos, C., Serandour, A.A., Ali, H.R., Brown, G.D., Atkins, A., Rueda, O.M., Holmes, K.A., Theodorou, V., Robinson, J.L.L., Zwart, W., Saadi, A., Ross-Innes, C.S., Chin, S.-F., Menon, S., Stingl, J., Palmieri, C., Caldas, C., Carroll, J.S., 2013. Endogenous Purification Reveals GREB1 as a Key Estrogen Receptor Regulatory Factor. Cell Rep. 3, 342–349. 10.1016/j.celrep.2013.01.010

Moran, K.M., Delville, Y., 2024. A hamster model for stress-induced weight gain. Horm. Behav. 160, 105488. 10.1016/j.yhbeh.2024.105488

Pałucha-Poniewiera, A., Podkowa, K., Rafało-Ulińska, A., Brański, P., Burnat, G., 2020. The influence of the duration of chronic unpredictable mild stress on the behavioural responses of C57BL/6J mice. Behav. Pharmacol. 31, 574–582. 10.1097/FBP.0000000000000564

Payne, J.L., 2003. The role of estrogen in mood disorders in women. Int. Rev. Psychiatry 15, 280–290. 10.1080/0954026031000136893

Perez-Villalba, A., Teruel-Martí, V., Ruiz-Torner, A., Olucha-Bordonau, F., 2005. The effect of long context exposure on cued conditioning and c-fos expression in the rat forebrain. Behav. Brain Res. 161, 263–275. 10.1016/j.bbr.2005.02.028

Porcelli, A.J., Delgado, M.R., 2017. Stress and decision making: effects on valuation, learning, and risk-taking. Curr. Opin. Behav. Sci. 14, 33–39. 10.1016/j.cobeha.2016.11.015

R Core Team, 2014. R: A language and environment for statistical computing. R Foundation for Statistical Computing.

Rabasa, C., Dickson, S.L., 2016. Impact of stress on metabolism and energy balance. Curr. Opin. Behav. Sci., Diet, behavior and brain function 9, 71–77. 10.1016/j.cobeha.2016.01.011

Reed, J.C., 1998. Bcl-2 family proteins. Oncogene 17, 3225–3236. 10.1038/sj.onc.1202591

Robbins, J., 2001. KCNQ potassium channels: physiology, pathophysiology, and pharmacology. Pharmacol. Ther. 90, 1–19. 10.1016/S0163-7258(01)00116-4

Saha, S., Datta, K., Rangarajan, P., 2007. Characterization of mouse neuronal Ca2+/calmodulin kinase II inhibitor α. Brain Res. 1148, 38–42. 10.1016/j.brainres.2007.02.018

Schoenebeck, B., Bader, V., Zhu, X.R., Schmitz, B., Lübbert, H., Stichel, C.C., 2005. Sgk1, a cell survival response in neurodegenerative diseases. Mol. Cell. Neurosci. 30, 249–264. 10.1016/j.mcn.2005.07.017

Simpkiss, J., Devine, D., 2003. Responses of the HPA axis after chronic variable stress: Effects of novel and familiar stressors. Neuro Endocrinol. Lett. 24, 97–103.

Süss, H., Willi, J., Grub, J., Ehlert, U., 2021. Estradiol and progesterone as resilience markers? – Findings from the Swiss Perimenopause Study. Psychoneuroendocrinology 127, 105177. 10.1016/j.psyneuen.2021.105177

Takahashi, A., Chung, J.-R., Zhang, S., Zhang, H., Grossman, Y., Aleyasin, H., Flanigan, M.E., Pfau, M.L., Menard, C., Dumitriu, D., Hodes, G.E., McEwen, B.S., Nestler, E.J., Han, M.-H., Russo, S.J., 2017. Establishment of a repeated social defeat stress model in female mice. Sci. Rep. 7, 12838. 10.1038/s41598-017-12811-8

Van Den Bos, R., Harteveld, M., Stoop, H., 2009. Stress and decision-making in humans: Performance is related to cortisol reactivity, albeit differently in men and women. Psychoneuroendocrinology 34, 1449–1458. 10.1016/j.psyneuen.2009.04.016

Watanabe, Y., Gould, E., McEwen, B.S., 1992. Stress induces atrophy of apical dendrites of hippocampal CA3 pyramidal neurons. Brain Res. 588, 341–345. 10.1016/0006-8993(92)91597-8

Wei, R., Li, D., Jia, S., Chen, Y., Wang, J., 2023. MC4R in Central and Peripheral Systems. Adv. Biol. 7, 2300035. 10.1002/adbi.202300035

Wickham, H., Averick, M., Bryan, J., Chang, W., McGowan, L., François, R., Grolemund, G., Hayes, A., Henry, L., Hester, J., Kuhn, M., Pedersen, T., Miller, E., Bache, S., Müller, K., Ooms, J., Robinson, D., Seidel, D., Spinu, V., Takahashi, K., Vaughan, D., Wilke, C., Woo, K., Yutani, H., 2019. Welcome to the Tidyverse. J. Open Source Softw. 4, 1686. 10.21105/joss.01686

Wolfes, A.C., Dean, C., 2020. The diversity of synaptotagmin isoforms. Curr. Opin. Neurobiol. 63, 198–209. 10.1016/j.conb.2020.04.006

Woolley, C.S., Gould, E., McEwen, B.S., 1990. Exposure to excess glucocorticoids alters dendritic morphology of adult hippocampal pyramidal neurons. Brain Res. 531, 225–231. 10.1016/0006-8993(90)90778-A

Wu, T., Hu, E., Xu, S., Chen, M., Guo, P., Dai, Z., Feng, T., Zhou, L., Tang, W., Zhan, L., Fu, X., Liu, S., Bo, X., Yu, G., 2021. clusterProfiler 4.0: A universal enrichment tool for interpreting omics data. The Innovation 2. 10.1016/j.xinn.2021.100141

Xie, M., Xiong, Y., Wang, H., 2024. The regulative role and mechanism of BNST in anxiety disorder. Front. Psychiatry 15. 10.3389/fpsyt.2024.1437476

Yohn, C.N., Ashamalla, S.A., Bokka, L., Gergues, M.M., Garino, A., Samuels, B.A., 2019. Social instability is an effective chronic stress paradigm for both male and female mice. Neuropharmacology 160, 107780. 10.1016/j.neuropharm.2019.107780

Zhang, X., Pöschel, B., Faul, C., Upreti, C., Stanton, P.K., Mundel, P., 2013. Essential Role for Synaptopodin in Dendritic Spine Plasticity of the Developing Hippocampus. J. Neurosci. 33, 12510–12518. 10.1523/JNEUROSCI.2983-12.2013

Zou, Y.-F., Wang, F., Feng, X.-L., Li, W.-F., Tian, Y.-H., Tao, J.-H., Pan, F.-M., Huang, F., 2012. Association of DRD2 gene polymorphisms with mood disorders: A meta-analysis. J. Affect. Disord. 136, 229–237. 10.1016/j.jad.2010.11.012

